# Systematic evaluation of chromosome conformation capture assays

**DOI:** 10.1101/2020.12.26.424448

**Authors:** Betul Akgol Oksuz, Liyan Yang, Sameer Abraham, Sergey V. Venev, Nils Krietenstein, Krishna Mohan Parsi, Hakan Ozadam, Marlies E. Oomen, Ankita Nand, Hui Mao, Ryan MJ Genga, Rene Maehr, Oliver J. Rando, Leonid A. Mirny, Johan Harmen Gibcus, Job Dekker

**Author notes:** contributed equally (Betul Akgol Oksuz, Liyan Yang).

## Abstract

Chromosome conformation capture (3C)-based assays are used to map chromatin interactions genome-wide. Quantitative analyses of chromatin interaction maps can lead to insights into the spatial organization of chromosomes and the mechanisms by which they fold. A number of protocols such as in situ Hi-C and Micro-C are now widely used and these differ in key experimental parameters including cross-linking chemistry and chromatin fragmentation strategy. To understand how the choice of experimental protocol determines the ability to detect and quantify aspects of chromosome folding we have performed a systematic evaluation of experimental parameters of 3C-based protocols. We find that different protocols capture different 3D genome features with different efficiencies. First, the use of cross-linkers such as DSG in addition to formaldehyde improves signal-to-noise allowing detection of thousands of additional loops and strengthens the compartment signal. Second, fragmenting chromatin to the level of nucleosomes using MNase allows detection of more loops. On the other hand, protocols that generate larger multi-kb fragments produce stronger compartmentalization signals. We confirmed our results for multiple cell types and cell cycle stages. We find that cell type-specific quantitative differences in chromosome folding are not detected or underestimated by some protocols. Based on these insights we developed Hi-C 3.0, a single protocol that can be used to both efficiently detect chromatin loops and to quantify compartmentalization. Finally, this study produced ultra-deeply sequenced reference interaction maps using conventional Hi-C, Micro-C and Hi-C 3.0 for commonly used cell lines in the 4D Nucleome Project.

## Introduction

Chromosome conformation capture (3C)-based assays (1) have become widely used to generate genome-wide chromatin interaction maps (2). Analysis of chromatin interaction maps has led to detection of several features of the folded genome. Such features include precise looping interactions (0.1-1 Mb scale) between pairs of specific sites that appear as local dots in interaction maps. Many of such dots represent loops formed by cohesin-mediated loop extrusion that is stalled at convergent CTCF sites (3–5). Loop extrusion will also produce other features in interaction maps including stripe-like patterns anchored at specific sites that block loop extrusion, and the effective depletion of interaction across such blocking sites leading to domain boundaries (insulation). At the Mb scale interaction maps of many organisms including mammals display checkerboard patterns that represent the spatial compartmentalization of two main types of chromatin: active and open A-type chromatin domains and inactive and more closed B-type chromatin domains (6). Finally, analysis of chromatin interaction frequencies and their dependence on genomic distance has been used to test mechanistic models of chromosome folding (5–10).

The Hi-C protocol has evolved over the years. While initial protocols used restriction enzymes such as HindIII that produces relatively large fragments of several kb (6), over the last 5 years Hi-C using DpnII or MboI digestion has become the protocol of choice for mapping chromatin interactions at kilobase resolution (3). More recently, Micro-C, which uses MNase instead of restriction enzymes as well as a different cross-linking protocol, was shown to allow generation of nucleosome-level interaction maps for yeast and several human and mouse cell lines (11–13). It is critical to ascertain how key parameters of these 3C-based methods, including cross-linking and chromatin fragmentation, quantitatively influence the detection of chromatin interaction frequencies and the detection of different chromosome folding features that range from local looping between small cis elements to global compartmentalization of Mb-sized domains. Such benchmarking of methods for detection of chromosome folding is an important goal of the 4D Nucleome Project (14). Here we systematically assessed how different cross-linking and fragmentation methods yield quantitatively different chromatin interaction maps. We identified optimal protocol variants for either loop or compartment detection and then used this information to develop a new Hi-C protocol (Hi-C 3.0) that can detect both loops and compartments relatively efficiently. In addition to providing benchmarked protocols, this work also produced important resource datasets including ultra-deep chromatin interaction maps using Micro-C, conventional Hi-C and Hi-C 3.0 for key cell lines widely used by the 4D Nucleome project.

## Results

We set out to explore how two key parameters of 3C-based protocols, cross-linking and chromatin fragmentation, determine the ability to quantitatively detect chromatin compartment domains and loops. We selected three cross-linking chemistries widely used to cross-link chromatin: 1) 1% formaldehyde, conventional for most 3C-based protocols, 2) 1% FA followed by an incubation with 3 mM disuccinimidyl glutarate (FA+DSG protocol), and 3) 1% FA followed by an incubation with 3 mM Ethylene glycol bis(succinimidylsuccinate) (FA+EGS protocol) (Figure 1a). We selected 4 different nucleases for chromatin fragmentation: MNase, DdeI, DpnII and HindIII, which fragment chromatin in sizes ranging from single nucleosomes to multiple kilobases. Combined, the 3 cross-linking and 4 fragmentation strategies yield a matrix of 12 distinct 3C-based protocols (Figure 1b). To determine how performance of these protocols varies for different states of chromatin we applied this matrix of protocols to multiple cell types and cell cycle stages. We analyzed 4 different cell types: pluripotent H1-hESCs, differentiated endoderm (DE) cells derived from H1-hESCs, fully differentiated Human Foreskin Fibroblasts (HFF, and a clonal derivate HFFc6) and HeLa S3 cells. Furthermore, we analyzed two cell cycle stages: G1 and mitotic HeLa S3 cells (Figure 1).

**Figure 1:**
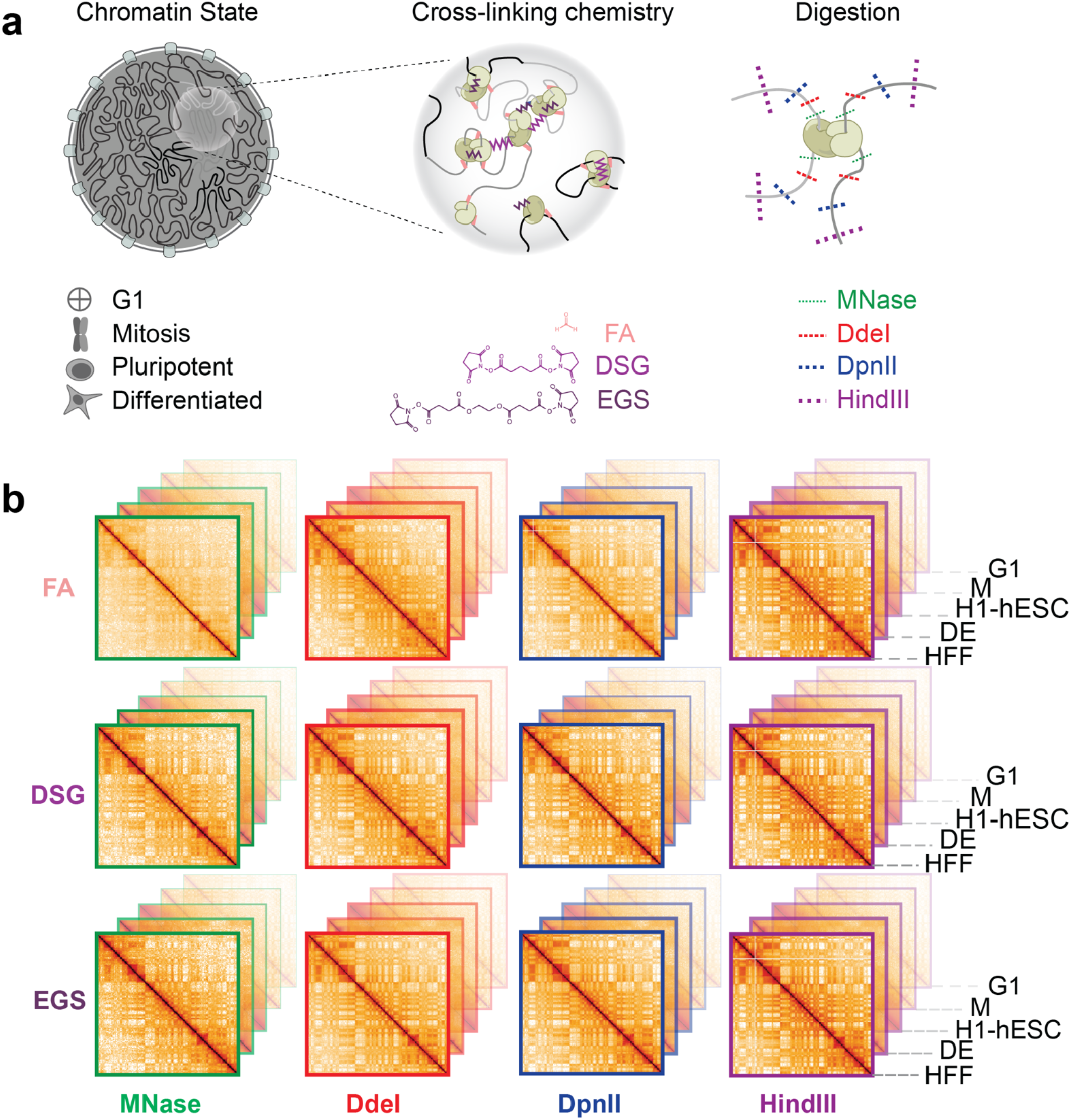
Outline of the experimental design. a. Experimental design for conformation capture on cells with the indicated chromatin states (left), using various cross-linking chemistries (middle) and digestion methods (right). b. Representation of interaction maps generated by combinations of experimental conditions depicted in panel a.

We first assessed the size range of the chromatin fragments produced after digestion by the twelve protocols for HFF cells (see Methods). Digestion with HindIII resulted in 5-20 kb DNA fragments; DpnII and DdeI produced fragments of 0.5-5kb; and MNase digested up to the level of mononucleosomes (~150 bp) (Supplemental Figure 1). Different cross-linking chemistries did not affect the size ranges produced by the different nucleases much, though DSG cross-linking lowered digestion efficiency slightly (Supplemental Figure 1b).

We applied this matrix of 3C-based protocols to the different cell types and cell cycle stages to produce a total of 63 libraries (12 protocols for H1-hESCs, DE and HFF cells; 9 protocols for each of the HeLa S3 cell cycle states, G1, M and non-synchronous cells (NS)); Supplemental Table 1). These libraries were then sequenced each on a single lane of a HiSeq4000 instrument, producing ~150-200 million uniquely mapping read pairs for each experiment (Supplemental Table 1).

### All tested 3C-based protocols can differentiate between cell states

We first assessed global correlations between the 63 datasets. We applied HiCRep to calculate all pairwise correlations between the datasets (15, 16) and then performed hierarchical clustering (Supplementary Figure 1c). The datasets cluster primarily by cell type and state, indicating that all protocols detect cell state-dependent differences. All datasets obtained with HeLa S3 G1 and NS cells are highly correlated (Supplemental Figure 1d). This is probably due to the fact that most cells in a NS cell population are in interphase. All datasets obtained with H1-hESC and DE cells are also highly correlated, reflecting their very close biological relationship (Supplemental Figure 1f). Further, mitotic datasets are the most distinct from all other datasets independent of the experimental protocol (Supplemental Figure 1e). Overall, protocols that use MNase to digest the chromatin have lower correlations with other protocols for a given cell type and cell state (Supplemental Figure 1d-g).

### Additional cross-linking produces more intra-chromosomal interactions in all cell states

Since chromosomes occupy individual territories, intra-chromosomal (cis) interactions are more frequent than inter-chromosomal (trans) interactions (17). The ratio of the number of interactions found in cis and trans is commonly used as indicator of Hi-C library quality given that inter-chromosomal interactions are a mixture of true chromatin interactions and interactions that are the result of random ligations (see below) (17, 18). For all enzymes and cell types tested, we found that the addition of DSG or EGS to the standard FA cross-linking decreased the percentage of trans interactions in favor of cis interactions genome-wide (Figure 2a (HFF); Supplemental Fig.2a-e (H1-hESC, DE, Hela S3)).

**Figure 2:**
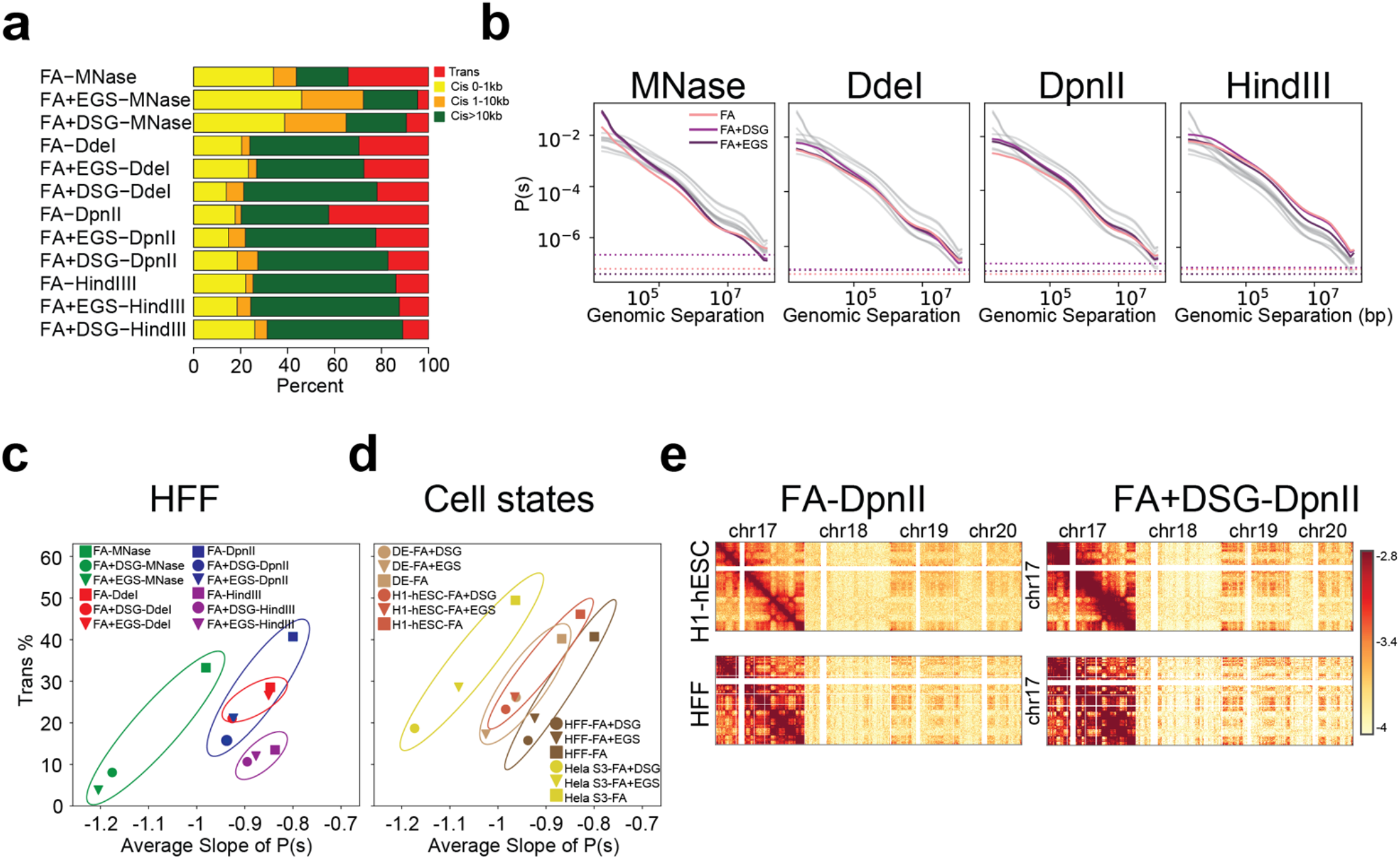
Distance dependent interaction frequency and the number of inter-chromosomal interactions change across protocols that use various enzyme and cross-linker combinations. Figures a, b and c created using 12 protocols that are performed in HFF cells. a. The number of valid pairs in each of the 12 protocols categorized by genomic distance. b. Distance dependent contact probability as detected by the set of 12 protocols split by used nucleases. Gray lines indicate all datasets, colored lines indicate data obtained with the indicated nuclease. c. The relationship between the percentage of trans interactions and the average slope of the distance dependent contact probability separated by cross-linker and enzyme combinations. Oval lines group datasets obtained with the same nuclease. d. The relationship between percentage of trans interactions and average slope of the distance dependent contact probability separated by cell type. Only experiments in which chromatin cross-linked with FA, FA+DSG or FA+EGS and digested with DpnII are shown. e. Interaction map of chromosome 17 with chromosomes 17, 18, 19 and 20 for FA or FA+DSG cross-linking and DpnII digestion, in H1-hESC and HFF. Total trans interactions for FA-DpnII protocols in H1-hESC: 47.7%, HFF: 42.5% and for FA+DSG-DpnII protocols in H1-hESC: 25% and HFF: 17.3%.

With respect to intra-chromosomal interactions, we noticed two distinct patterns. First, digestion into smaller fragments resulted in a relative increase in short range interactions. Nucleosome level digestion resulted in a higher number of interactions between loci separated by less than 10 kb, whereas digestion with either DdeI, DpnII or HindIII resulted in a relatively larger number of interactions between loci separated by more than 10 kb (Figure 2a). Distance dependent contact probability (*P*(*s*)) plots (6, 7) reflected these distributions (Figure 2b (HFF)). Second, visual inspection of *P*(*s*) plots showed that the addition of either DSG or EGS resulted in a steeper decay in interaction frequency as a function of genomic distance for all fragmentation protocols. By plotting the percentage of trans interactions as a function of the average slope of *P*(*s*), Figure 2c illustrates the main findings regarding the effects of fragmentation, cross-linking strategies and their interplay for HFF cells: for a given chromatin fragmentation level, additional cross-linking with DSG or EGS reduces the percentage of trans interactions, likely because spurious ligations are reduced (see below), and steepens the slope of *P*(*s*). Further, the smaller the chromatin fragments produced, the steeper the decay of *P*(*s*).

This relationship between percentage of trans interactions and the slope of the *P*(s) plots was observed for all cell types (Figure 2d, Supplemental Figure 2a-c) and cell cycle stages studied here (Supplemental Figure 2d, 2e). We note that the different cell types and cell cycle stages differed in the range of slopes. For instance, for a given fragmentation level, interaction frequencies obtained with non-synchronized HeLa S3 cells showed steeper slopes than those observed with H1-hESCs, DE and HFF cells (Figure 2d). This was the case for all protocols, indicating a biological difference between these cell lines.

One of the sources of noise in 3C-based experiments is the result of random ligation events between freely moving fragments that did not become cross-linked and now can ligate to any other such free fragments. This specific source of noise will mostly produce trans interactions, and can also constitute a noticeable proportion of very long range cis interactions where true signal is low. Differences in the percentage of trans interactions detected with different protocols can in part be the result of different levels of random ligation events. To test this we ascertained the frequency with which we observed interactions between the mitochondrial and nuclear genomes, as these are considered to form only through random ligation (Supplemental Figure 2f). We could not use this noise metric for experiments using MNase because MNase completely degrades the mitochondrial genome. Interestingly we observed that random ligations between genomic and mitochondrial DNA are the lowest when chromatin was fragmented with HindIII, and generally higher when chromatin was fragmented in smaller segments with DpnII or DdeI. The use of additional DSG or EGS cross-linking reduced random ligation in experiments using DpnII or DdeI, but did not further reduce random ligations when HindIII was used. The reduced noise in experiments using HindIII or in experiments using DpnII with additional cross-linkers is readily visible in chromatin interaction maps: for these protocols we observed a general decrease in trans contacts, while uncovering a stronger inter-chromosomal compartmental pattern (Figure 2e). We conclude that noise as a result of random ligation can be reduced by either using 6 bp-cutters (HindIII) or, when using more frequent cutters (DpnII, DdeI), by using additional DSG or EGS cross-linking. The reduced noise improves trans compartment detection and possibly long range cis interactions.

### Quantitative detection of compartmentalization is enhanced by long fragments and extra cross-linkers

To investigate compartmentalization we determined the positions of A and B compartments by eigenvector decomposition (19). The eigenvector associated with the largest eigenvalue typically represents the checkerboard pattern associated with compartments (6, 19). We analyzed the effect of the experimental protocol on compartment domain detection and on quantification of the preference with which A domains interact with A domains, and B domains with B domains (compartmentalization strength). We did this analysis for all cell states except for mitotic cells that do not display compartmentalized chromosomes (7). Correlation between compartment profiles of all experiments showed that the greatest difference in profiles can be attributed to cell type (Supplemental Figure 3a). Within each cell type, positions of compartment domains obtained with different protocols were highly similar (Spearman correlation >0.8). These high correlations indicate that compartment definition *per se,* and the genomic positions of compartmental domains, are largely insensitive to the experimental protocol (Supplemental Figure 3a).

We used the contrast of the checkerboard patterns in contact matrices to quantify the strength of the compartmentalization. Visual inspection of interaction matrices (binned at 100 kb resolution) suggested that the contrast between the domains that make up the checkerboard pattern can vary between protocols. For instance, for HFF cells cross-linked with only FA, interaction matrices obtained from MNase digestion displayed a relatively weak checkerboard pattern, whereas those obtained with HindIII digestion showed much stronger patterns (Fig 3a).

**Figure 3:**
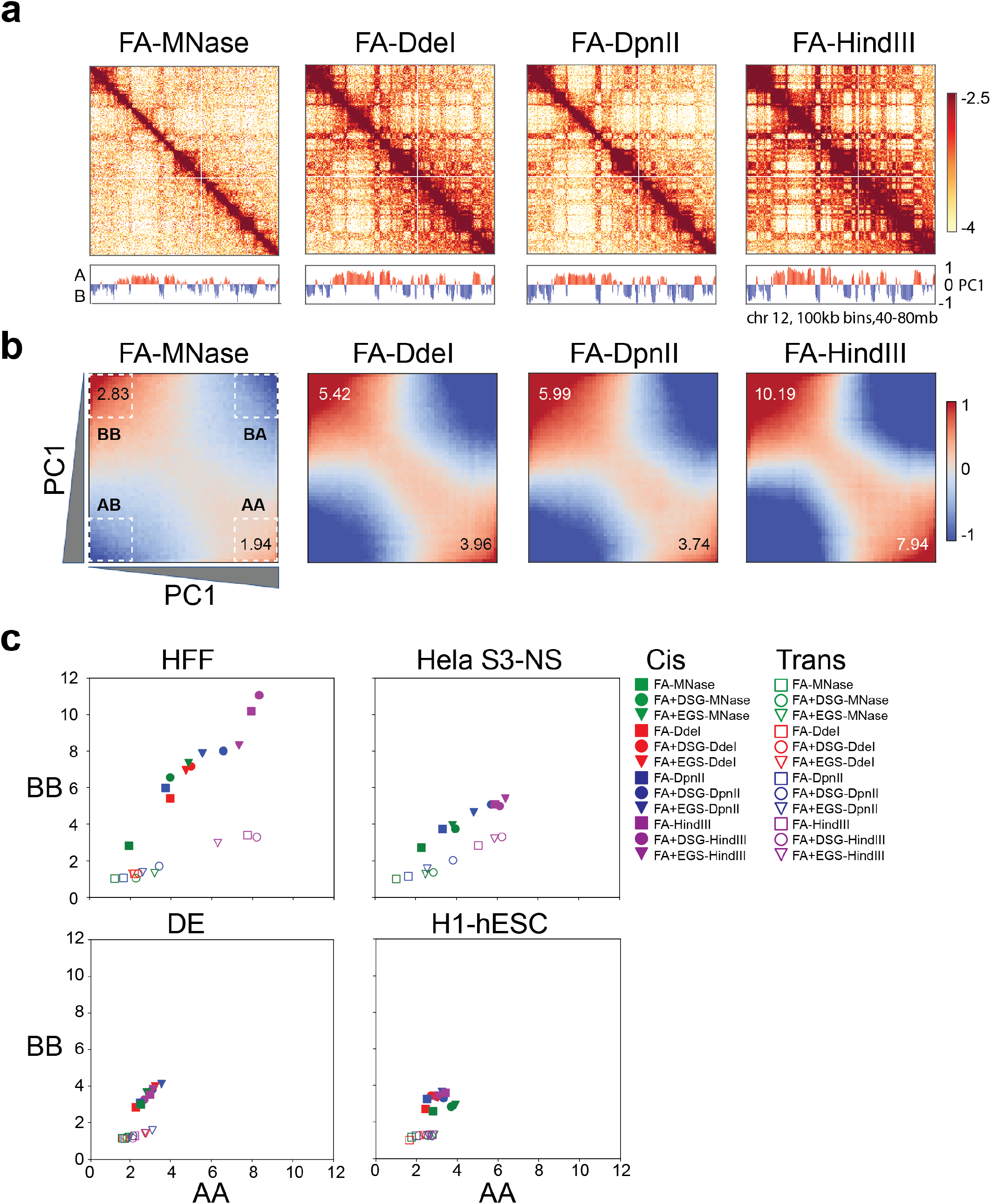
Cross-linking the chromatin with DSG or EGS and digesting it with HindIII strengthen compartment signals. a. Interaction maps for HFF cells obtained with experiments where the chromatin is cross-linked with FA only and digested with MNase, DdeI, DpnII and HindIII, respectively. PC1 values of the genomic region are displayed below the figure. b. Saddle plots of the genome-wide interaction maps of the data shown in Figure 3a. A-A and B-B compartment signals in cis get stronger with increasing fragment size. c. Quantification of the compartment strength using saddle plots of cis and trans interactions for 12 protocols applied to HFF cells, 9 protocols to Hela S3 NS, 12 protocols to DE, 12 protocols to H1-hESC. Y-axis represents the quantification for the strongest 20% of B-B and x-axis represents the quantification of the strongest 20% of A-A interactions.

The contrast of the checkerboard pattern can be quantified as follows: genomic loci are ranked by their eigenvector values, normalized for their expected interaction frequencies and then the interactions between them are plotted to produce a “saddle plot” (19). Preferential A-A and B-B interactions are quantified by dividing the A-A interactions and B-B interactions of the top 20% of the strongest A and B loci (based on their eigenvector values), by the sum of the corresponding A-B interactions (Figure 3b).

Saddle plot analysis (ranked eigenvector values; see Methods) revealed three trends in the detected compartment strength. First, protocols that generate larger fragments (e.g. using HindIII; Figure 3b, 3c) and protocols that include additional DSG or EGS cross-linking produced quantitatively stronger compartment patterns (Figure 3c; Supplemental Figure 3b, 3c, 3d, 3e). This trend held for all 4 cell types analyzed here. Second, different cell types differed in compartment strength: HFF cells displayed the strongest compartment pattern, while H1-hESCs displayed weak compartments regardless of the capture protocol used. Third, compartment strength was much stronger in cis than in trans.

Furthermore, some protocols, including the conventional Hi-C protocol (cross-linked with FA and digested with DpnII) and MNase-based protocols (Micro-C, regardless of cross-linking protocol) did not detect preferential B-B interactions in trans (Supplemental Figure 3), whereas such preferred B-B interactions are detected when Hi-C is performed with HindIII (Supplemental Figure 3d, 3e)).

Additionally, trans preferential A-A interactions are more frequent than trans preferential B-B interactions for all protocols and cell types. In summary, detected compartment strength is stronger, both in cis and in trans, when using protocols that produce larger fragments or employ additional cross-linking.

### Chromatin loops are better detected in experiments with fine fragmentation and additional DSG cross-linking

Of all structural Hi-C features, the detection of loops depends the most on sequencing depth. We applied 1) conventional Hi-C using FA and DpnII digestion (FA-DpnII); 2) Hi-C using DSG in addition to FA cross-linking and DpnII digestion (FA+DSG-DpnII); and 3) the standard Micro-C protocol (FA+DSG-MNase) to two cell types, H1-hESC and HFFc6, and sequenced the resulting libraries to a depth of 2.4-3.9 billion valid interactions (combined two biological replicates). HFFc6 cells are a subclone of HFF cells that are commonly used by 4D Nucleome Consortium (14). Interaction maps of data obtained from these “deep” datasets showed quantitative differences in interactions for both H1-hESC and HFFc6 (Supplemental Figure 4a-4b). As compared to conventional Hi-C (FA-DpnII), the use of additional cross-linking with DSG and finer fragmentation produces contact maps with more contrast and more pronounced focal enrichment of specific looping contacts. We then used a reimplementation of the HICCUPS approach to identify looping interactions that appear as dots in deep contact maps using settings that accurately reproduce previously published loops in GM12878 cells (3, 12) (see Methods).

In H1-hESCs we detected 3,951 loops with the FA-DpnII protocol, 12,396 loops with the FA+DSG-DpnII protocol and 22,507 loops with the FA+DSG-MNase protocol (Supplemental Figure 4c). For HFFc6 these numbers were 13,867, 22,934 and 36,988 respectively (Figure 4a). Additional DSG cross-linking allowed detection of many thousands more loops. Fragmenting chromatin with MNase further increased the number of loops that were detected. (Figure 4a, Supplemental Figure 4c). Here, detectability is defined as the ability of the dot calling algorithm (using published parameters) to distinguish a point-like region on a heatmap as having a statistically significant increase in contacts relative to its local surroundings. We then extracted all possible subsets of detected looping interactions from overlapping two or more called loop lists in order to investigate variations in the properties of these interactions across the subsets (Figure 4a). While a large fraction of loops was detected by all three protocols, we found that the protocols with additional cross-linking (FA+DSG-DpnII) and finer fragmentation (FA+DSG-MNase) detected a large set of additional loops (Fig. 4a).

**Figure 4.**
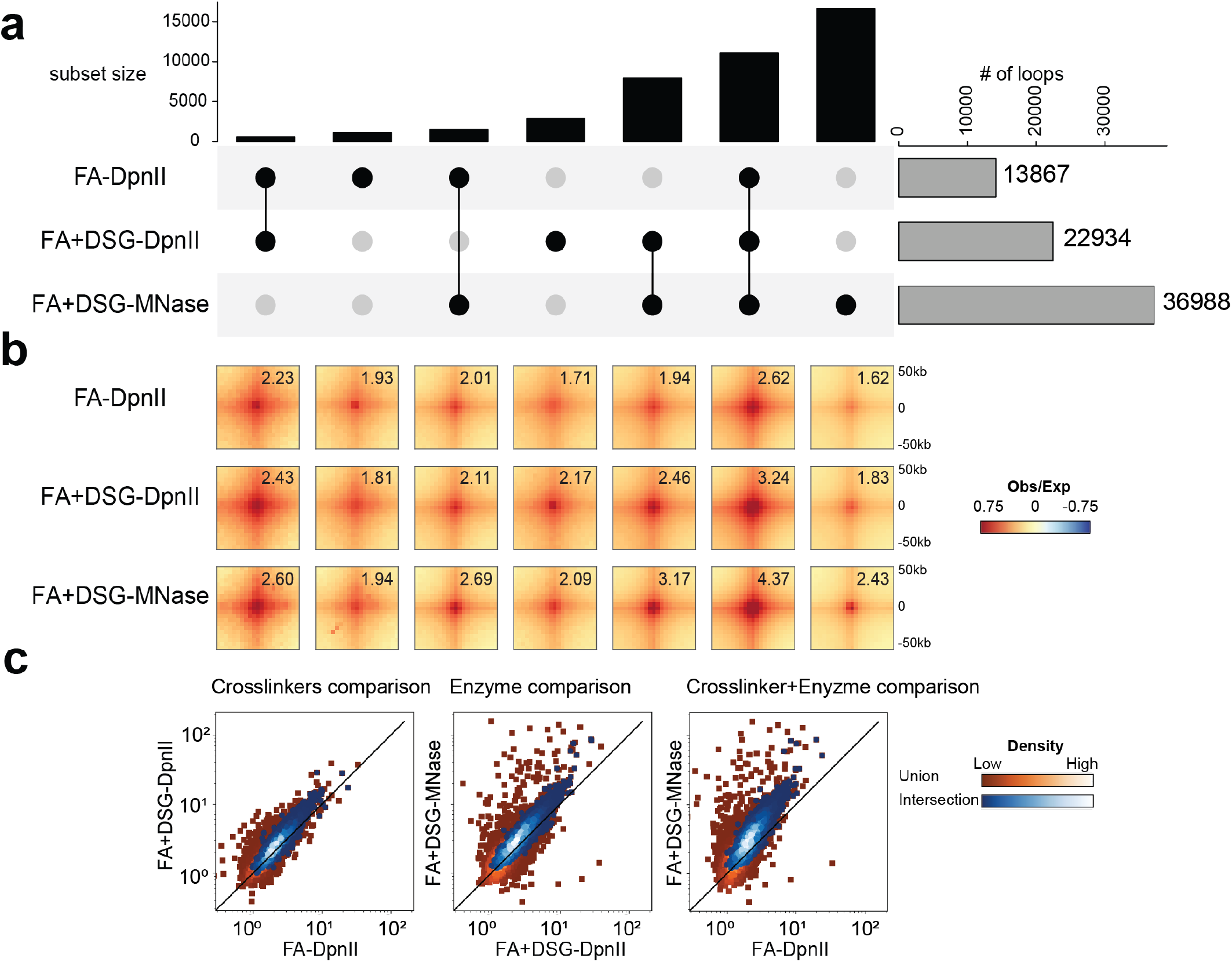
Chromatin loops are better detected in experiments with fine fragmentation and DSG cross-linking. a. Upset plot of loops detected in protocols performed using HFFc6 shows the 1) total number of loops detected in FA-DpnII, FA+DSG-DpnII and FA+DSG-MNase on the right side (gray bars), 2) number of loops detected in one, two or three of these protocols shown in black bars. Loops found with only one or multiple protocols are highlighted and connected with black dots. b. The pileups of the loops in HFFc6 for every set of loops shown in Figure 4a. Numbers in each pileup represent the signal enrichment at the loop compared to local background. See methods for quantification of loop strength. c. Scatter plots to investigate the relative strengths of individual loops between pairs of protocols for HFFc6 cells. The pairs are (from left to right): FA-DpnII v/s FA+DSG-DpnII (different crosslinking with the same enzyme), FA+DSG-DpnII v/s FA+DSG-MNase (different enzymes with same crosslinking) and FA-DpnII v/s FA+DSG-MNase (different enzymes and different crosslinking). Loop strengths were calculated the same as in panel d but for individual looping interactions. The scatters are done for two sets of looping interactions - the union of all locations from the three protocols (red squares) and interaction of all locations from the three protocols (blue circles). The color scale represents the density of loop interactions present at those strengths.

When we aggregated interaction data for the various subsets of loops detected with one or multiple different protocols, we observed a focal increase in interaction frequency for all subsets of loops for all datasets; even for data obtained with protocols where that subset of loops was not detected as significantly enriched (Figure 4b for HFFc6, Supplemental Fig. 4d for H1-ESC). For instance, loops only detected with the FA+DSG-MNase protocol were also visible in aggregated data obtained with the FA-DpnII protocol. Quantifying the strength of the different subsets of loops detected by one or multiple protocols, we found that loops detected by all three protocols were the strongest, while loops detected only by the FA+DSG-MNase protocol were relatively weak.

We then defined a consensus set of loops that are called in all three deep datasets and analyzed this set using interaction datasets obtained with the matrix of 12 protocols described in Figure 1 that differ in cross-linking and fragmentation strategies. We observed a gradual increase in average loop strength as the fragment sizes were smaller and extra DSG or EGS cross-linkers were used (Supplemental Figure 4f, 4g).

To explore this in another way, we quantified the individual strengths of each of the loops in the set of consensus loops. The majority of loops were strengthened by addition of DSG as an extra cross-linker (Figure 4c left panel). Further, the loop strength increased by digestion with MNase as compared to DpnII (Figure 4c, middle). We conclude that the use of additional cross-linkers and enzymes that fragment chromatin in smaller fragments independently contribute to the loop strength and the number of loops that was detectable. Loops were strongest when both additional cross-linkers were used and chromatin was fragmented with MNase (Figure 4c, right plot). Similar results were obtained when we used the union of loops detected in all three deep datasets (Figure 4c, Supplemental Figure 4e).

A looping interaction is defined by a pair of interacting loci, anchors. When each anchor is involved in only 1 looping interaction, the number anchors will be twice the number of loops. In contrast, when anchors engage in multiple looping interactions with other anchors, the number of anchors will be smaller than twofold the number of loops (5). To examine this, we plotted the number of anchors as a function of the number of loops for the sets of loops detected in the deep datasets. For HFFc6 cells we observed that for loops detected with the FA-DpnII protocol the number of anchors was close to twice the number of loops (Figure 5a). Interestingly, for protocols that used additional cross-linkers (FA-DSG-DpnII) and finer fragmentation (FA-DSG-MNase), where we had detected increased numbers of loops (above), the increase in the number of additional anchors is less than twice the increase in the number of additional loops. This indicates that many of the newly identified loops involved anchors that were also detected with FA-DpnII (Figure 5a, Supplemental Figure 5a). In other words, many of additionally detected loops are arranged along stripes emanating from the same anchors.

**Figure 5:**
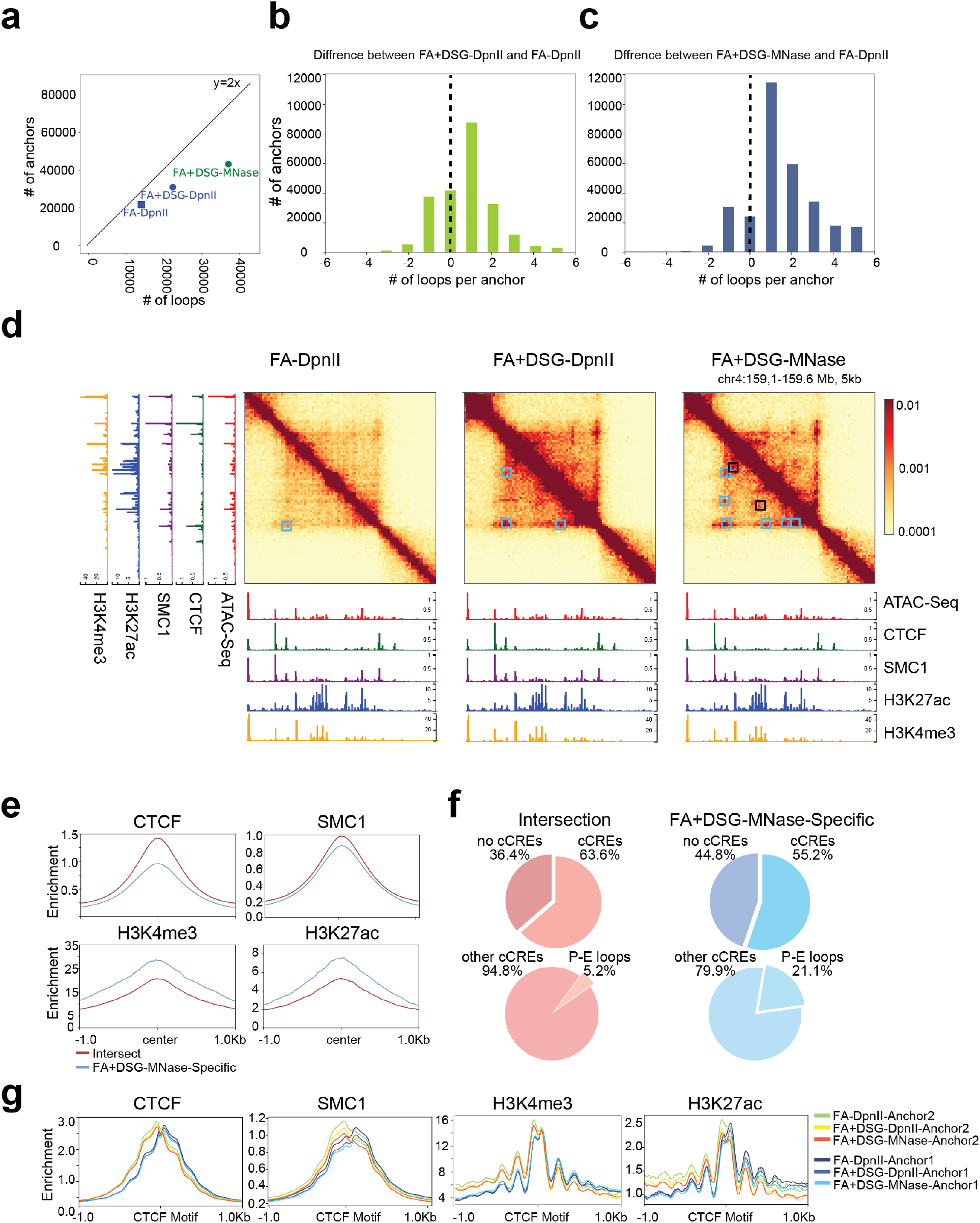
Characterization of interactions and chromatin features of loop anchors detected with different protocols. a. The number of loops detected in HFFc6 (x-axis) plotted against the number of loop anchors (y-axis). For y=2x depicts the expected relationship when each anchor is engaged in only 1 loop. b,c. The number of FA-DpnII loops subtracted from the number of FA+DSG-DpnII (b) or FA+DSG-MNase (c) loops detected at the same anchors; the union list of the of the plotted protocols was used. d. Interaction maps for 3 protocols applied to HFFc6 cells and CUT$Run/CUT&Tag data for CTCF, SMC1, H3K4me3 and H3K27ac. Cyan squares highlight the loop anchors detected with all three protocols. Black squares indicate loop anchors detected with FA+DSG-MNase only (20). e. CTCF, SMC1, H3K4me3 and H3K27ac enrichments at loop anchors detected by all protocols (union) or FA+DSG-MNase alone in HFFc6. Open chromatin regions within anchor coordinates were used to center average enrichments. f. Candidate Cis Regulatory elements (cCREs) detected in common and FA+DSG-MNase specific loop anchors from Figure 5e (top) and stratified percentage of Promoter-Enhancer cCREs without CTCF enrichment (bottom). g. Enrichment of CTCF, SMC1, H3K4me3 and H3K27ac separated between left (Anchor1) and right (Anchor 2) anchor for loop anchors detected in HFFc6 using FA-DpnII, FA+DSG-DpnII or FA+DSG-MNase.

To further investigate this we directly determined the number of loops that a given anchor engaged in as detected by different protocols and then calculated the difference between them. For each anchor, we subtracted the number of loops detected by the FA-DpnII protocol from the number of loops detected using the FA+DSG-DpnII or the FA-DSG-MNase protocol. We found that using extra cross-linkers (FA-DSG-DpnII) as well as finer fragmentation (FA-DSG-MNase) increased the number of detectable loops for most anchors (Figure 5b, 5c, Supplemental Figure 5b, 5c). We conclude that protocols that use additional cross-linkers and finer fragmentation detect more loops in two ways: first, more loops are detected per anchor, and second, additional looping anchors are detected.

We split loop anchors into two categories: 1) anchors detected with more than 1 protocol and 2) anchors detected with only 1 protocol. We observed that anchors detected with at least 2 protocols were engaged in multiple loops (loop “valency” >1). In contrast, anchors that were detected in only 1 protocol mostly had a loop valency of 1 (Supplemental Figure 5d, 5e). Interestingly, for H1-ESCs the majority of additional loops detected with FA-DSG-MNase protocol (62%) involve two anchors not detected with other protocols. In comparison for HFFc6 this was only 21% indicating that most new loops shared at least one anchor with loops detected with other protocols.

We investigated factor binding (CTCF and cohesin (SMC1), YY1 and RNA polII) and chromatin state (H3K4Me3, H3K27Ac) at the two categories of loop anchors. We used publicly available datasets (21,22) and new data generated using a variety of techniques (Cut&Run (23), Cut&Tag (24), ChIP Seq and ATAC-Seq (25)) for this analysis. Examples are shown in Figure 5d. Some loop anchors are detected in all protocols and in the example shown these correspond to sites bound by CTCF and cohesin (cyan squares). Other loop anchors are only detected with the FA-DSG-MNase protocol (black squares). In this example these do not correspond to sites by CTCF and cohesin, but are enriched in H3K27Ac and H3K4Me3. Possibly, the ability of different protocols to detect various loop anchors is related to factor binding and chromatin state. To investigate this genome-wide we aggregated CTCF, cohesin, YY1, RNA PolII binding data and histone modification data (H3K4me3 and H3K27ac) at loop anchors detected with all protocols or with only FA-DSG-MNase (Figure 5e). Interestingly, in HFFc6 cells we found that FA+DSG-MNase-specific loop anchors are less enriched for CTCF and SMC1 but more enriched for H3K4me3 and H3K27ac compared to the loop anchors that are detected by all three protocols which are more enriched for CTCF and SMC1 but less enriched for H3K4me3 and H3K27ac (Figure 5e, Supplemental Figure 5g).

Next, we examined the predicted cis regulatory elements that are located at shared loop anchors across the three deep datasets obtained with the three protocols and at loop anchors detected only with the FA+DSG-MNase protocol. We used candidate cis-Regulatory element (cCREs) predictions from ENCODE (26) which uses DNase hypersensitive regions, CTCF, H3K4me3 and H3K27ac ChIP seq data for predicting and annotating cCREs with and without CTCF binding sites. We found that the majority of the shared anchors have cCREs but only a small part of these cCREs are predicted promoter or enhancer elements without CTCF site (5.2% for HFFc6, 9.8% for H1-ESC; Figure 5f, Supplemental Figure 5g). In contrast, around half of the FA-DSG-MNase-specific anchors have predicted cCREs and for this subset the number of predicted promoter or enhancer elements without CTCF site is higher compared to loop anchors detected with all protocols (21% for HFFc6, 30% for H1-ESC; Figure 5f, Supplemental Figure 5g).

Finally, we compared the chromatin organization at CTCF-enriched loop anchors with respect to the orientation of the CTCF binding motif. Remarkably, using Cut&Tag or Cut&Run data we found an asymmetric distribution of signal for all factors (Figure 5g), including CTCF (Cut&Tag data). Both CTCF and cohesin signals are skewed towards the inside of the loop. We noted that the Cut&Tag data was generated with an antibody against the N-terminus of CTCF (Figure 5g). We also analyzed Cut&Run data that was generated with an antibody directed against the C-terminus of CTCF (Supplemental Figure 5h) and observed signal enrichment skewed at CTCF sites towards the outside of the loop. These observations are consistent with the orientation of CTCF binding to its motif and interactions between the N-terminus of CTCF with cohesion on the inside of the loop (27). The stronger enrichment of H3K4Me3 and K3K27Ac on the inside of the loop is intriguing, but the mechanism of this asymmetry is not known.

### Insulation quantification is robust to experimental variations

Next we investigated chromatin insulation, i.e. the reduced interaction probability across loop anchors and domain boundaries (28–30). We used the insulation metric that quantifies the frequency of interactions across any genomic locus within a set window size. We quantified insulation for each 10 kb bin by aggregating interactions across each bin over a 200 kb window (31). Local minima in this metric represent positions of insulation where interactions between domains are relatively infrequent. The local depth of the minimum is a measure for the strength of the insulation (31).

First, we compared the insulation strength as detected with the deep datasets obtained with the FA-DpnII, FA+DSG-DpnII and FA+DSG-MNase protocols in HFFc6. We observed that the distribution of the insulation strengths is bimodal: for each dataset we identify a relatively large set of very weak boundaries, and a smaller set of strong boundaries (Supplemental Fig. 6a). Insulation at the weak boundaries is very small, and possibly noise (Supplemental Fig. 6d). We focus on the set of strong boundaries. We aggregated insulation profiles at 1) loop anchors detected with each of the three deep datasets (Supplemental Figure 6b (left)), 2) strong insulation boundaries (Supplemental Figure 6b (middle)) and 3) loop anchors that are at strong insulation boundaries (Supplemental Figure 6b (right)). We find that insulation at these elements is very similar for each of the three deep datasets, indicating that the different protocols perform comparably in quantitative detection of insulation. We note that in general insulation at strong boundaries is stronger than at loop anchors, possibly because of our stringent threshold for boundary detection.

Second, we investigated whether insulation strength depends on sequencing depth. We compared two biological replicates; one with ~150 M interactions (matrix data, Supplemental Figure 6c (left)) and the other with 2.5 B interactions (deep data, Supplemental Figure 6a) for data obtained with the FA-DpnII, FA+DSG-DpnII and FA+DSG-MNase protocols. Deeper sequencing reduced the relative number of weak boundaries, suggesting these are due to noise (Supplemental Figure 6a, 6c (left)). The majority (> 85 %) of strong boundaries are detected in both deep data and the less deeply sequenced data obtained with the matrix of 12 protocols, and the insulation scores of these shared strong boundaries are highly correlated across all datasets (r > 0.80) (Supplemental Figure 6c (right)).

Third, we investigated the number and the strength of the boundaries detected using data obtained with the matrix of 12 protocols for HFF, H1-hESC, DE cells and the 9 protocols for HelaS3. The number of strong boundaries detected with protocols using MNase fragmentation was between 7,000-8,000 compared to other protocols which identified between 5,000-6,300 boundaries, irrespective of cross-linking chemistry. Insulation strength at boundaries detected with each protocol was very similar (Supplemental Figure 6d (right)). We observed the same results for H1-hESC (Supplemental Figure 6e-h).

We find a positive correlation between boundary strength and the number of protocols that detect that boundary (Supplemental Figure 6i, 6j). Focusing on the set of boundaries that are detected by at least half of the protocols we then investigated how insulation varied for data obtained with the matrix of 12 protocols. We found that insulation strength is very similar for data obtained with all protocols (Supplemental Figure 6k). Similarly, when insulation is aggregated at the set of loop anchors detected by all three deep datasets using Hi-C data obtained with the matrix of 12 protocols we detect only minor variations in insulation. In summary insulation detection and quantification was robust to variations in protocol (Supplemental Figure 6l).

### Double digestion with both DdeI and DpnII improves loop detection, while maintaining ability to quantitatively detect compartments

Our comprehensive comparison of protocols showed that compartment signals were strongest for datasets obtained with protocols that generate longer fragments (i.e. HindIII) or that employ additional cross-linkers (FA+DSG and FA+EGS). On the other hand, looping interaction strength was superior for datasets obtained with protocols that generate small fragments (MNase) and employ additional cross-linkers. We considered whether a single protocol could be designed that optimally captures both compartments and loops. To this end, we tested the effect of double digestion with DdeI and DpnII after cross-linking with FA+DSG (FA+DSG-DdeI+DpnII, referred to as “Hi-C 3.0”). As expected, we observed that using two enzymes further shortened the fragment size compared to individual enzyme digestion (Supplemental Figure 7a, 7b). Applying this protocol to HFFc6 cells, we generated two deeply sequenced biological replicates (3.3 billion valid interactions combined). For comparison, we also generated deeply sequenced dataset from HFFc6 cells using only DdeI digestion (FA-DSG-DdeI; 2.7 billion valid interactions combined) in addition to the deeply sequenced libraries digested with only DpnII or MNase described above.

We found that the FA+DSG-DdeI+DpnII protocol affected the distance dependent contact probability (Figure 6a). Compared to data obtained by single DdeI or DpnII digests, contacts increased for loci separated by less than 10 kb, making the results from this protocol more similar to results obtained with protocols using MNase digestion. Yet, longer distance contacts resembled data obtained with protocols using single restriction enzymes more than data obtained with protocols using MNase. Combined, this protocol improved short range signal without loss of long-range signal (Figure 6b).

**Figure 6 :**
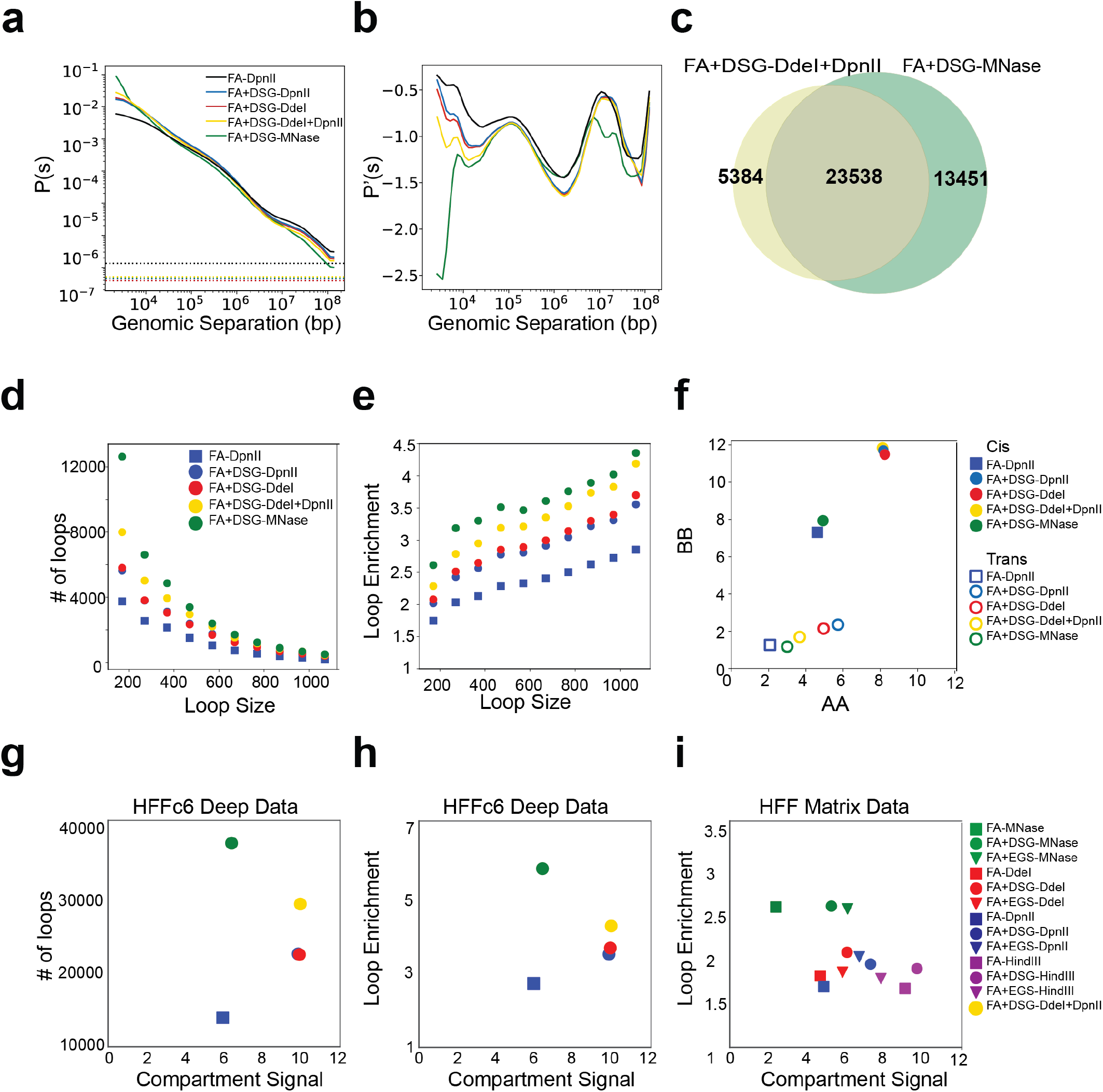
Loop detectability and strength increase when the chromatin is digested with two restriction enzymes while preserving strong compartment signal. a. *P*(*s*) plot showing distance dependent contact probability of interactions detected with 5 protocols applied to HFFc6 cells. Dashed lines show the percentage of trans interactions for each dataset. b. Derivative of the *P*(s) plots shown in panel a. c. Venn diagram shows the overlap between the number of loops detected with FA+DAG-DdeI+DpnII and FA+DSG-MNase. d. The number of loops detected within 100kb intervals (loop size) starting at 70kb. Bin intervals: 70-170 kb, 170-270 kb, 270-370,……,970-1070 kb. e. Loop strength of 1,000 loops sampled from 100 kb intervals (Figure 6d). When less than 1,000 loops were available, loop strengths for available loops were used. f. A-A (x-axis) and B-B (y-axis) compartment strengths in cis and trans derived from saddle plot analysis. g. The relationship between the number of loops and compartment strength for 5 protocols applied to HFFc6 cells. h. Compartment strength (x-axis) compared to loop enrichment quantification for 10,000 loops sampled from HFFc6 deep data (2,000 loops were sampled from each of 5 protocols). i. Compartment strength obtained from the matrix of 12 protocols applied to HFF (x-axis) compared to loop enrichment for the set of 10,000 loops sampled from the deep data (panel h) using interaction data obtained from the same matrix of 12 protocols (y-axis).

When considering the reproducibility of loop calls between the double digest and MNase libraries, we found that the majority of the loops detected with FA+DSG-DdeI+DpnII overlapped with those detected with FA+DSG-MNase (Figure 6c). Consistent with our previous observations, loop anchors detected in both protocols had stronger CTCF and cohesin enrichment, whereas protocol specific anchors (for both FA+DSG-DdeI+DpnII and FA+DSG-MNase) were more enriched for H3K4me3 and H3K27ac (Supplemental Figure 7c). We found that the smaller fragments due to the double digestion reveal more pronounced looping interactions compared to single restriction enzyme digestion. The loop calling algorithm applied to this interaction map detected ~6,000 more looping interactions than what was discovered with data generated with either single DpnII or single DdeI digestion (Figure 6d). Furthermore, the average enrichment of contacts at these looping interactions was also higher for data obtained with the double digestion protocol. Nonetheless, the MNase library remained superior in detecting loops - both in number and in contact enrichment (Figure 6e). Importantly, when we investigated compartmental interactions we found that the shortening of restriction fragments did not result in a loss of quantitative detection of preferential compartmental interactions (Figure 6f). In conclusion, the FA+DSG-DdeI+DpnII double digest protocol allows for the efficient detection of both loops and compartments in single protocol (Figure 6g-6h).

## Discussion

We present a systematic evaluation of experimental parameters of 3C-based protocols that determine the ability to detect and quantify aspects of chromosome folding. Fragmentation level and cross-linking chemistry determine the ability to detect chromatin loops or compartmentalization in different ways. Loop detection improves when chromatin is cross-linked with additional (DSG) cross-linking and cut into small fragments. Loops detected with such protocols are more enriched for cis elements like enhancers and promoters as compared to sets of loops detected with conventional Hi-C. However, this comes at a cost of a reduced ability to quantitatively detect compartmentalization in cis and in trans. Quantification of compartmentalization improves with longer fragments such as those produced with DpnII in conventional Hi-C. Compartment strength improves further with additional cross-linkers or when chromatin is digested in even longer fragments, e.g. using HindIII. A protocol using two restriction enzymes (DpnII and DdeI) and additional DSG cross-linking (Hi-C 3.0) combines strengths of the MNase-based Micro-C protocol to detect loops and Hi-C protocols to detect stronger compartments.

Fragmentation level and cross-linking chemistry determine assay performance by affecting the level of noise due to random ligation events in datasets (17). We find that smaller fragments result in more random ligation events. Possibly the number of cross-links per fragment is low for small fragments, leading to a higher mobility during the assay. These mobile fragments ligate to other such fragments randomly, which will mostly increase very long-range and inter-chromosomal interactions. Consistent with this interpretation is that random ligation events diminish when additional cross-linking is used or when chromatin is fragmented into larger fragments. This results in a decrease in inter-chromosomal interactions and steeper *P*(*s*) plots. Improved signal to noise ratios allowed better detection of loops, compartments, and more meaningful inter-chromosomal interactions.

Detection of compartmentalization strength is improved when protocols are used that produce relatively long fragments and include additional cross-linking. Possibly, compartmental interactions are more difficult to capture than looping interactions that are closely held together by cohesin complexes. Recently, we found that interfaces between compartment domains appear relatively unmixed (32). Longer fragments or extra cross-linkers may be required to more efficiently capture contacts across these interfaces. Interestingly, cell type-specific differences in strength of compartmentalization are only observed with some protocols. Conventional Hi-C (FA+DpnII) suggests that compartmentalization strength is quite similar in H1-ESCs, Hela S3, DE cells and HFF. However, when Hi-C is performed with additional cross-linkers and/or with restriction enzymes that produce longer fragments, compartmentalization in HFF and Hela S3 are much stronger, while compartmentalization strength for H1-ESCs and DE are unaffected. This suggests that quantitative differences in cell type-specific chromosome organization can be missed or underestimated depending on the 3C-based protocol.

Insights into the influence of experimental parameters of chromatin interaction data led us to test a single Hi-C protocol, that we refer to as Hi-C 3.0, that can be used for better detection of both loops and compartments. This protocol employs double cross-linking with FA and DSG followed by double digestion with two restriction enzymes (DpnII and DdeI). This produces shorter fragments than those in conventional Hi-C, but not as short as in the nucleosome-sized fragments in Micro-C. This protocol allowed detection of many thousands more loops than conventional Hi-C (FA+DpnII), while also detecting strong compartmentalization. Therefore we propose that Hi-C 3.0 would be the Hi-C method of choice for widespread chromatin interaction studies.

The very deeply sequenced Hi-C, Micro-C and Hi-C 3.0 datasets we produced for H1-ESCs,, and HFFc6 cells will be useful resources for the chromosome folding community given that these cell lines are widely used for method benchmarking and analysis by the 4D nucleome project (14). Further, the comprehensive collection of chromatin interaction data generated with the matrix of twelve 3C-based protocol variants for each cell line can also be a valuable resource for benchmarking computational methods for data analysis given their different cross-linking distances and chemistry, fragment lengths and noise levels.

## Acknowledgements

This work was supported by a grant from the National Institutes of Health Common Fund 4D Nucleome Program to J.D. and L.A.M (U54-DK107980), a grant from the National Human Genome Research Institute (NHGRI) to J.D. (HG003143). J.D. is an investigator of the Howard Hughes Medical Institute. N.K. was supported by Human Frontiers Science Program (HFSP) grant LT000631/2017-L. We thank Drs. Stephen Henikoff and Derek Janssenss for sharing Cut&Run data.

## Author contributions

J.D and J.H.G. conceived the study. L.Y and N.K performed 3C-based assays. J.H.G. coordinated data collection. B.A.O. led all data analysis and performed data analysis. S.A. performed data analysis. S.V.V. contributed analysis tools and performed analysis. H.O. pre-processed the sequencing data. A.N., H.O. and J.H.G. designed the cLIMS system to store metadata, K.M.P and R.M. differentiated H1-hESC to DE and cultured the cells, M.E.O performed mitotic data synchronization and HFFc6 ATAC-seq experiments. H.M. and R.MJ.G generated H1-hESC ATAC-seq and HFFc6 Cut&Tag datasets. O.J.R. and L.A.M. contributed data analysis. All authors contributed to writing the manuscript.

## Data availability

Data are available at GEO under accession number GSE163666. Supplemental table 1 list datasets accessible through the 4DN data portal includes 4DN accession numbers.

## Code availability

Scripts and notebooks that are used in this manuscripts are located here: https://github.com/dekkerlab/matrix_paper

## Methods

### cLIMS: A Laboratory Information Management System for C-Data

cLims is a web-based lab information management system tailored for chromosome conformation capture experiments. It can be used to organize, store and export metadata of various experiment types such as HiC, 5C, ATAC-Seq, etc. The metadata organization is compatible with 4DN DCIC standards and data to cLIMS can be used to export 4DN DCIC and GEO systems with one click.

For the matrix project, we had increasing levels of detail in metadata, growing number of experiments, long time periods between data creation and submission and many people working on the same data sets, hence cLIMS helped us to keep this information properly maintained. The details included cell line, assay, treatments, sequencing, contributor’s information. This will also help us in reproducibility of experiments.

cLIMS has been developed using the Django web framework on the back-end and HTML5 and Javascript libraries on the front-end. It is running on PostreSQL database and Apache web server and can be hosted on major Linux distributions.

### Cell line culture and fixation

#### HFFc6

HFFc6 was cultured according to 4DN SOP (https://data.4dnucleome.org/biosources/4DNSRC6ZVYVP/). Cells were grown at 37°C under 5% CO2 in 75cm^2^ flasks containing Dulbeco’s Modified Eagle Medium (DMEM), supplemented with 20%, heat inactivated Fetal Bovine Serum (FBS). For sub-culture, cells were rinsed with 1x DPBS and detached using 0.05% trypsin at 37 °C for 2-3 minutes. Cells were typically split every 2-3 days at a 1:4 ratio and harvested while sub-confluent, ensuring they would not overgrow.

#### H1-hESC

Human Embryonic stem cells (H1 – WiCell, WA01, lot # WB35186) were cultured in mTeSR1 media (StemCell Technologies, 85850) under feeder-free conditions on Matrigel H1-hESC-qualified matrix (Corning, 354277, lot # 6011002) coated plates at 37°C and 5% CO_2_. H1 cells were daily fed with fresh mTeSR1 media and passaged every 4-5 days using ReLeSR reagent (StemCell Technologies, 05872). Cells were dissociated into single cells with TrypLE Express (Thermo Fisher, 12604013).

#### Fixation protocol

Final harvest of 5 million HFFc6 and H1-hESC cells was performed after washing twice with Hank’s Buffered Salt Solution (HBSS) before cross-linking in HBSS with 1% Formaldehyde for 10 minutes at room temperature. Formaldehyde was quenched with glycine (128 mM final concentration) at room temperature for 5 minutes and on ice for an additional 15 minutes. Cells were washed twice with DPBS before pelleting and flash freezing with liquid nitrogen into 5 million aliquots. Alternatively, formaldehyde fixed cells were centrifuged at 800x*g* and subjected to additional cross-linking with either 3mM Disuccinimidyl glutarate (DSG) or Ethylene glycol bis (succinimidylsuccinate) (EGS), freshly prepared and diluted from a 300mM stock in DMSO, for 40 minutes at room temperature. DSG and EGS cross-linked cells were both quenched with 0.4M glycine for 5 minutes and washed twice with DPBS, supplemented with 0.5% Bovine Serum Albumin, before flash freezing with liquid nitrogen into 5 million aliquots.

#### Hi-C protocol

Chromosome conformation capture was performed as described previously and we refer to Belaghzal et al. (33) for a step-by-step version similar to this protocol. Briefly, 5×10^6^cross-linked cells were lysed for 15 minutes in ice cold lysis buffer (10 mM Tris-HCl pH8.0, 10 mM NaCl, 0.2% Igepal CA-630) in the presence of Halt protease inhibitors (Thermo Fisher, 78429). Then, the cells were disrupted by homogenization with pestle A for 2x 30 strokes. An aliquot if 8 μL was taken to later assess chromatin integrity. Remaining chromatin was solubilized in 0.1% SDS at 65°C for 10 minutes, quenched by 1% Triton X-100 (Sigma, 93443) and digested with 400 units of either HindIII (R0104 in NEBuffer 2.1), DdeI (R0175, in NEBuffer 3.1) or DpnII (R0543 in NEBuffer 3.1) for 16 hours at 37°C. Samples were incubated at 65 ̊C for 20 minutes to inactivate the restriction enzyme after which 10 μL was set aside to assess digestion efficiency. Fill-in of digested overhangs by DNA polymerase I, large Klenow fragment (NEB, M0210) in the presence of 250 nM biotin-14-dCTP (HindIII; Thermo Fisher, 19518018) or biotin-14-dATP (DdeI, DpnII; Thermo Fisher, 19524016) for 4 hours at 23°C was performed prior to ligation with 50 μL T4 DNA ligase (Thermo Fisher, 15224090) for 4 hours at 16°C in a total volume of 1.2 mL. Cross-links of ligated chromatin were reversed at 65°C overnight by 2 separate 50 μL additions of 10 mg/mL proteinase K (Fisher, BP1750I-400). DNA was isolated by adding 2.6 mL saturated phenol pH 8.0: chloroform (1:1) to 1.3 mL of sample. The mixture was vortexed and spun down in phase-lock tubes (Quiagen, 129065) before standard precipitation with 100% ethanol in the presence of 1/10 vol/vol of 3 M sodium acetate pH 5.2. DNA cleanup and desalting was performed using an AMICON Ultra Centrifuge filter, following manufacturer’s instructions (EMD Millipore, UFC5030BK). RNA was removed by incubation with 1 μL of 1 mg/mL RNAase A for 30 minutes at 37°C in a total of 100 μL TLE (10 mM Tris-HCl, 0.1 mM EDTA in milliQ) and quantified on a 0.7% agarose gel. Biotin was removed at 20°C for 4 hours in a 50 μL reaction for every 5 μg of DNA using 15 units of T4 DNA polymerase (NEB, M0203L) and 25 nM dATP and 25nM dGTP in NEBuffer 3.1 (no dTTP and dCTP). Polymerase was inactivated for 20 mins at 75°C and placed at 4°C. Volume was brought up to 130 μL and DNA was sheared for 3 minutes using a Covaris sonicator (E220 evolution: Duty Cycle 10%, Intensity 5, Cycles per Burst 200 or M220: Peak Incident Power 50W, Duty Cycle 20%, Cycles per Burst 200) and size selected with Agencourt AMPure® XP (Beckman Coulter, A63881) to obtain 150 - 350 basepair fragments, validated by DNA gel electrophoresis. DNA was repaired by adding a cocktail of 20 μL of the end-repair mix [3.5X NEB ligation buffer (NEB, B0202S), 17.5 mM dNTP mix, 7.5 units of T4 DNA polymerase (NEB, M0203L), 25 units of T4 polynucleotide kinase (NEB, M0201S), 2.5 units Klenow polymerase Polymerase I (NEB, M0210L)] to 50μL DNA solution at 20°C for 30 minutes, followed by a 20 minute incubation at 75°C to inactivate Klenow polymerase. For every library, at least 10 μL of streptavidin coated Dynabeads™ MyOne™ Streptavidin C1 (Thermo Fisher, 65001) in LoBind tubes (Eppendorf, 022431021) were prepared by washing the beads twice with Tris Wash Buffer [5 mM Tris-HCl pH8.0, 0.5 mM EDTA, 1 M NaCl, 0.05% Tween20] and resuspending in 400 μL of 2X Binding Buffer [10 mM Tris-HCl pH 8, 1 mM EDTA, 2 M NaCl]. The washed beads were added to the biotinylated DNA (brought to 400 μL TLE) and incubated for 15 minutes at room temperature under rotation. Thereafter, the beads were first washed with 1x Binding Buffer and then with TLE before final elution in 41μL TLE on a magnetic stand. Then, a 9 μL A-tailing mix consisting of 5 μL 10x NEBuffer 2.1, 1 μL of 10 mM dATP and 15 units of Klenow 3’ → 5’ exo- (NEB M0212L) was added to blunted ends and incubated at 37°C for 30 minutes, followed by inactivation for 20 minutes at 65°C. Beads were reclaimed, washed with 1x ligation buffer (from 5x T4 DNA ligase buffer, Thermo Fisher, 46300-018) and Illumina paired-end adapters were added by ligation with T4 DNA ligase (Thermo Fisher, 15224090) for 2 hours at room temperature. To determine the minimal number of PCR cycles needed to generate a Hi-C library, a PCR titration was performed prior to the production PCR (using Illumina primers PE1.0 and PE2.0). Primers were separated from the final library by size selection with AMpure XP (1:1 ratio) prior to 50 bp paired-end sequencing on an Illumina HiSeq 4000 sequencer (Thermo Fisher).

For each deep library repeat, we generated 4 Hi-C libraries in parallel (20× 10^6^ cells total) and sequenced each of the generated libraries on a single lane of an Illumina HiSeq 4000 flow cell.

#### Micro-C-XL protocol

The Micro-C XL protocol was adopted from Hsieh et al. and Krietenstein et al. (11, 12). Frozen cells were resupended in 200 ul cold 1x PBS (10 nM Na2HPO4, KH2PO4, pH 7.4, 137 nM NaCl, 2.7 nM KCl) per 1 million cells and split into 1 million cell aliquots. Note, 1x BSA (NEB, B9000S) was added to PBS prior resuspension and washing to reduce stickiness of HFFc6 cells to the tub walls. After 20 min incubation on ice, cells were collected by centrifugation (5000x *g* for 5 minutes), washed with 500 μL buffer MB#1 (10 mM Tris-HCl, pH 7.5, 50 nM NaCl, 5 mM MgCl2, 1 mM CaCl2, 0.2% NP-40, 1x Roche complete EDTA-free (Roche diagnostics, 04693132001), collected by centrifugation (5000x *g* for 5 minutes), and resuspended in 200 μL MB#1. Chromatin was incubated with MNase for 20 min at 37°C. MNase concentrations were chosen to yield mostly mono-nucleosomal fragments, as tested in prior digestion tests, typically 5-20 units MNase (Wortington Biochem, LS004798). The digestion was stopped by addition of 0.5 mM EGTA (Bioworld, 05200081) to a 1.5 mM final concentration and incubation at 65°C for 10 minutes. Chromatin aliquots were pooled for further processing. Here, the equivalent of 2.5 million cells input yielded the best results, more than 5 million cell-equivalent per aliquot is not recommended. The chromatin was collected by centrifugation (5000x *g* for 5 minutes), washed with 500 μL 1x NEBuffer 2.1 (NEB, B7202S), collected by centrifugation (5000x *g* for 5 minutes), and resuspended in 45 μL NEBuffer 2.1. DNA ends were dephosphorylated by addition of 5 μL rSAP (NEB, M0203) and incubation at 37°C for 45 min. The reaction was stopped by incubation at 65°C for 5 min. 5’ overhangs were generated by 3’ resection. Here, 40 μL pre-mix (5 μL 10x NEBuffer 2.1, 2 μL 100 mM ATP (Thermo Fisher, R0441), 3 μL 100 mM DTT, 30 μL H2O) and 8 μL Large Klenow Fragment (NEB, M0210L) and 2 μL T4 PNK (NEB, M0201L) were added to the sample in respective order. The reaction was incubated at 37°C for 15 minutes. The DNA overhangs were filled with biotinylated nucleotides by addition of 100 μL pre-mix (25 μL 0.4 mM Biotin-dATP (Thermo Fisher, 19524016), 25 μL 0.4 mM Biotin-dCTP (Thermo Fisher, 19518018), 2 μL 10 mM dGTP and 10 mM dTTP (stock solutions: NEB, N0446), 10 μL 10x T4 DNA Ligase Reaction Buffer (NEB B0202S), 0.5 μL 200x BSA (NEB, B9000S), 38.5 μL H2O) and incubation at 25°C for 45 minutes. The reaction was stopped by addition of 12 μL 0.5 M EDTA (Thermo Fisher, 15575038) and incubation at 65°C for 20 minutes. The chromatin was collected by centrifugation (10000x *g*), washed in 500 μL 1x Ligase Reaction Buffer, and collected by centrifugation (10000x *g*). The chromatin pellet was resuspended in 2500 μL ligation reaction buffer (1x NEB Ligase buffer, 1x NEB BSA, 12500 U NEB T4 Ligase (NEB, M0202L)) and incubated rotating at room temperature for 2.5-3 hours. After proximity ligation, the chromatin was collected and resuspended in 200 μL 1x NEBuffer 1 (NEB, B7001S) and 200 U NEB Exonuclease III (NEB, M0206S). For de-proteination and reverse cross-linking, 25 μL ProteinaseK (25mg/ml in TE with 50% glycerol) and 25 μL 10% SDS (Thermo Fisher, 15553035) were added and the sample was incubated at 65°C o/n. The DNA was first phenol:chloroform purified and second purified with DNA Clean & Concentrator Kit (Zymo, D4013). The 300 bp sized Micro-C library was purified via 1.5 % agarose gel electrophoresis and extracted with Zymoclean Gel DNA Recovery Kit (Zymo Research, D4002) with a final elution volume of 50 μL. 5 μL Dynabeads™ MyOne™ Streptavidin C1 beads (Thermo Fisher, 65001) were washed twice with 300 μL 1x TBW (5 mM Tris-HCl, pH 7.5, 0.5 mM EDTA, 1 M NaCl) and suspended in 150 μL 2x TBW (10 mM Tris-HCl, pH 7.5, 1 mM EDTA, 2 M NaCl). 100 μL H2O and 150 μL washed Streptavidin beads in 2x TBW were added to the sample and incubated rotation at room temperature for 20 minutes. The beads were washed twice with 300 μL 1x TBW and resuspended in 50 μL TE buffer. Sequencing libraries were prepared with NEBNext^®^ Ultra™ II DNA Library Prep Kit for Illumina^®^ (NEB, E7645) according to protocol, except for the DNA purification and size selection prior PCR. Here, adaptor-ligated DNA was pulled-down via bound Streptavidin beads and washed twice with 300 μL 1x TBW and once with 0.1x TE. Finally, the beads were resuspended in 20 μL 0.1x TE. PCR amplification, sample indexing, and DNA purification after PCR was performed according to (NEB, E7645) using NEBNext^®^ Multiplex Oligos for Illumina^®^. The samples were sequenced on an Illumina HiSeq 4000 on 50 base pair paired end mode.

#### Size range of chromatin fragments produced after digestion

Cells were cross-linked, lysed and digested as with the Hi-C protocol (see above). Then, cross-links were reversed and DNA was isolated as in Hi-C, but without ligation and biotin incorporation. DNA was loaded on an Advanced Analytical Fragment Analyzer (Agilent) for size range analysis and data was analyzed with PROsize3 software (Agilent). PROsize3 traces were exported separately as 4×8 bins (32 total) ranging from 40-500; 500-1300; 1300-8000 and 8000-100000 basepairs. Size ranges of potential restriction sites (hg38) were identified with cooltools genome digest (https://cooltools.readthedocs.io/en/latest/cli.html?highlight=enzyme#cooltools-genome-digest).

#### Cut&Tag protocol

Samples were processed as previously described in Kaya-Okur et.al. (24), with few modifications. Briefly, approximately 100K cells per sample were permeabilized in the wash buffer (20 mM HEPES pH 7.5, 150 mM NaCl, 0.5 mM Spermidine, 1× Protease inhibitor cocktail) and then cells were coupled with activated concanavalin A-coated magnetic beads for 10 min at RT. Pelleted beads were resuspended in antibody buffer (Mix 8 μL 0.5 M EDTA and 6.7 μL 30% BSA with 2 mL Dig-wash buffer) with 1:100 dilution of SMC1 (Bethyl, cat# A300-055A) or CTCF antibody (Active motif, cat # 61311) and incubated overnight at 4 °C on a rotator. The next day, the pelleted bead complex was incubated with 1: 50 dilution of secondary antibody (guinea pig α-rabbit antibody, cat. # ABIN101961) in Dig-Wash buffer (20 mM HEPES pH 7.5, 150 mM NaCl, 0.5 mM Spermidine, 1× Protease inhibitor cocktail, 0.05% Digitonin) and incubated at RT for 30 min on rotator. After two washes in Dig-Wash buffer, 1:250 diluted pAG-Tn5 adapter complex in Dig-300 buffer (20 mM HEPES pH 7.5, 300 mM NaCl, 0.5 mM Spermidine, 1× Protease inhibitor cocktail, 0.05% Digitonin) were added to bead complex and incubated at RT for 1 hr. After two washes in Dig-300 buffer, beads were resuspended in 300 μL of Tagmentation buffer (20 mM HEPES pH 7.5, 300 mM NaCl, 0.5 mM Spermidine, 1× Protease inhibitor cocktail, 0.05% Digitonin,10 mM MgCl2) and incubated at 37 °C for 1 h 45 min. Samples were subjected to Proteinase K treatment and extracted tagmented DNA using Phenol:Chloroform:Isoamyl Alcohol (25:24:1). In preparation for Illumina sequencing, 21 μL DNA was mixed with 2 μL of a universal i5, 2 μL of a uniquely barcoded i7 primer, and 25 μL of NEBNext HiFi 2× PCR Master mix. The sample was placed in a thermocycler with a heated lid using the following cycling conditions: 72 °C for 5 min; 98 °C for 30 s; 14 cycles of 98 °C for 10 s and 63 °C for 30 s; final extension at 72 °C for 1 min and hold at 4 °C. Post-PCR clean-up was performed by adding 1.1× volume of Ampure XP beads and incubated for 15 min at RT, washed twice gently in 80% ethanol, and eluted in 30 μL 10 mM Tris pH 8.0. Final library samples were paired-end sequenced on Nextseq500.

#### Cut&Run Protocol

Cut&Run raw data (fastq files) of H1-hESC are downloaded from Janssens et al. 2018 (21) and raw files of HFFc6 are generated by Steve Henikoff Lab using Skene et al. 2017 protocol (23).

#### ATAC Seq Protocol

We have followed a published protocol to perform H1-hESC ATAC Seq experiments. The protocol details are described in Genga et al. 2019 (34).

ATAC-seq experiments on HFFc6 cells were performed following previously published protocol (Buenrostro, Wu, Chang, & Greenleaf, 2015) (25). Briefly, 50,000 cells per experiment were washed and lysed using a lysis buffer (0.1% NP-40, 10 mM Tris-HCl (pH 7.4), 10 mM NaCl and 3 mM MgCl2). Lysed cells were then transposed using the Nextera DNA library prep kit (Illumina #FC-121-1030) for 30 min at 37C, immediately followed by DNA collection using Qiagen MinElute columns (Qiagen #28004). Appropriate cycle numbers for amplification were determined for each sample individually using qPCR. Finally, primers were removed using AMpure XP beads (Beckman Coulter #A63881) prior to 2×50bp paired-end sequencing.

### Data analysis

#### Chromosome capture data processing

Distiller (https://github.com/mirnylab/distiller-nf) pipeline is used to process Hi-C and Micro-C datasets. First, sequencing reads were mapped to hg38 using bwa mem with flags-SP. Second, mapped reads were parsed and classified using the pairtools package (https://github.com/mirnylab/pairtools) to get 4DN-compliant pairs files. PCR and/optical duplicates removed by matching the positions of aligned reads with 2bp flexibility. Next, pairs were filtered using mapping quality scores (MAPQ > 30) on each side of aligned chimeric read, binned into multiple resolutions and low coverage bins are removed.

Finally multiresolution cooler files were created using the cooler package (https://www.biorxiv.org/content/10.1101/557660v1, https://github.com/mirnylab/cooler.git). We normalized contact matrices using the iterative correction procedure from Imakaev et al. 2012 (19). Interaction heatmaps were created using the “cooler show” command from the cooler package.

#### Hicrep correlations

We used HiCREP to do distance corrected correlations (15) of the various protocols and cell states. Correlation is calculated in two steps. First, interaction maps are stratified by genomic distances and the correlation coefficients are calculated for each distance separately. Second, the reproducibility is determined by a novel stratum-adjusted correlation coefficient statistic (SCC) by aggregating stratum-specific correlation coefficients using a weighted average.

#### Cis and Trans Ratio

Trans percent is calculated by dividing the total interactions between chromosomes with the sum of interactions within and between chromosomes (trans/cis+trans). Distance seperated cis interactions are calculated by dividing total interactions within specified distance of the chromosomes by the sum of interactions within and between chromosomes (cis of specific distance/cis+trans). Pairtools provides statistics for the numbers of interactions captured within and between chromosomes.

#### Scaling Plots

Scalings plots describe the decay of the average probability of contact between two regions on a chromosome as a function of the genomic separation between them.

As per best practices, scalings are typically computed for each chromosomal arm of the genome before being aggregated. In order to obtain the extent of each chromosomal arm, the sizes of the chromosomes and the positions of their associated centromeres must be obtained. The sizes of the chromosome were obtained using the *fetch_chromsizes* function that is found in the bioframe library (https://github.com/open2c/bioframe/blob/master/bioframe/io/resources.py#L61) and the starts and ends of the centromere were obtained from bioframe using *fetch_centromeres* (https://github.com/open2c/bioframe/blob/master/bioframe/io/resources.py#L109). The results of these two functions were combined to create a single list containing the extents of each chromosomal arm of the Human hg38 genome. For all libraries except those made from Hela S3 cells, all chromosome arms were used in the scaling calculation. For Hela libraries only chromosomes 4, 14, 17, 18, 20, and 21 were used.

We used the *diagram* function from the cooltools library (https://github.com/open2c/cooltools/blob/master/cooltools/expected.py#L541) to calculate scaling. This function takes in a cooler, extracts the table of non-zero read counts across the genome (known as the pixel table) and calculates the sum of read counts based on its distance from the main diagonal. It also simultaneously calculates the total number of possible counts obtainable at a given distance (called valid pairs) based on masking of region due to balancing and other use provided criteria. Additionally, this function also has the ability of transforming the read-counts obtained from the pixel table before aggregating the result. This is done by passing the appropriate use defined function to the “transforms” parameter of diagsum.

To obtain the scaling plots shown in the manuscript, for each library, the diagsum function was applied on the 1kb cooler associated with the library. 1kb is the recommended resolution to calculate scalings as it allows us to observe variations at the finest scales. Along with the cooler, the chromosomal arms extents were also provided using the regions argument. A transform (named “balanced”) was also applied to the data to convert raw read-counts to balanced read-counts. This was done by multiplying the count value with the associated row and column weights obtained from balancing the cooler.

The resulting output is a single table with 4 relevant columns: 1) “region” which describes what chromosome arm a specific row was obtained from; 2) “diag” which refers to the genomic separation at which the data was aggregated; 3) “balanced.sum” which is the sum of read-counts for that given region and genomic separation after they were transformed by the “balanced” transform and 4) “n_valid” the number of possible valid pairs at a given distance (as described earlier). The individual column values were aggregated over the different arms and then further aggregated into logarithmically spaced bins of genomic separation. Finally, the “balanced.sum” column was divided by the “n_valid” column to create the “balanced.avg” column that is a measure of the average number of contacts across the genomic for a given genomic separation. The curves shown in the main text are the “balanced.avg” values plotted as a function of “diag” for the different libraries.

In addition to the interaction decay within a chromosome, interaction between different chromosomes can also be quantified. This is done using the “*blocksum_asymm*” function in cooltools (https://github.com/open2c/cooltools/blob/master/cooltools/expected.py#L820) which uses a very similar methodology. Two sets of regions are provided to blocksum_asymm and “balanced.sum” and “n_valid” is calculated for every pair of regions (entire chromosomes in this case). Since the interactions are between two chromosomes there is no notion of genomic separation between two regions. The “balanced.avg” is calculated in the same manner as above and the mean of this value is visualized as horizontal dashed lines in the main text figures.

#### Average slope of scaling

In order to magnify small variations between the different libraries, we calculated “derivative curves” from the scaling curves. Derivative curves represent the rate of change of scaling curves as observed on a log-log scale. These are computed by taking the log of scaling data (both x and y), calculating the finite difference measure of the slope and smoothing that value with a gaussian kernel. The smoothing function used is *gaussian_filter1d* from the scipy library (with a spread of 1). The smooth finite difference values can be plotted as a function of distance as is the case for Fig 6b. Alternatively, the average value of this derivative is calculated and correlated with other features (as in Fig 2c,d)

#### Compartment Analysis

We assessed compartments using eigenvector decomposition on observed-over-expected contact maps at 100kb resolution separated for each chromosomal arm using the cooltools package derived scripts. Eigenvector that has the strongest correlation with gene density is selected, then A and B compartments were assigned based on the gene density profiles such that A compartment has high gene density and B compartment has low gene density profile (19). Spearman correlation (Supplementary Fig. 3a) was used to correlate the eigenvectors of different experiments performed with various protocols and cell states. Saddle plots were generated as follows: the interaction matrix of an experiment was sorted based on the eigenvector values from lowest to highest (B to A). Sorted maps were then normalized for their expected interaction frequencies; the upper left corner of the interaction matrix represents the strongest B-B interactions, lower right represents strongest A-A interactions, upper right and lower left are B-A and A-B respectively. To quantify saddle plots we took the strongest 20% of BB and strongest 20% of AA interactions and normalized them by the sum of AB and BA (top(AA)/(AB+BA) and top(BB)/(AB+BA)). Saddle quantifications were used to create the scatter plots in figure 3c and heatmaps in supplementary figure 3 that compare A and B compartments for all cell types. Both scatter plots and heatmaps in figure 3 and Supplementary figure 3 were created using the Matplotlib package from Python.

#### Identification of chromatin loops

The cooltools call-dots function (https://github.com/open2c/cooltools/blob/master/cooltools/cli/call_dots.py),a reimplementation of HICCUPS (3) was used to detect the chromatin loops that are reflected as dots in the interaction matrix. We used the following parameters to call the loops: fdr=0.1, diag_width=10000000, tile_size = 5000000, --max-nans-tolerated 4. We called dots in deep data at both 5kb and 10 kb resolutions, using MAPQ> 30 pairs and before merging the results using the criteria mentioned in Rao et al. 2014 (3). Briefly, to merge 5kb and 10 kb loop calls, both the reproducible 5kb calls and unique 10 kb calls were kept. Unique 5kb calls were kept if the genomic separation of the region was <100kb or if the dots were particularly strong (i.e.more than 100 raw interactions per 5kb pixel). More detailed explanations for dot calling can be found in Rao et al. 2014 and Krietenstein et al. 2020 (3, 12).

#### Comparison of loops detected in different protocols

*Bedtools intersect (35)* was re-implemented to overlap 2D loops between protocols. Since loop calls are fundamentally 2 dimensional data, they needed to be processed for use with bedtools (which operates on 1d data).

Each loop call consists of 6 coordinates: chrom1, start1, end1, chrom2, start2, end2. Since chrom1 is always the same as chrom2 for loop calls, we ignored these two columns and reduced our space to4 coordinates. Furthermore, to account for errors in the positioning of loop during the loop calling, we introduced the following margin of error around the called region (typically 10kb):

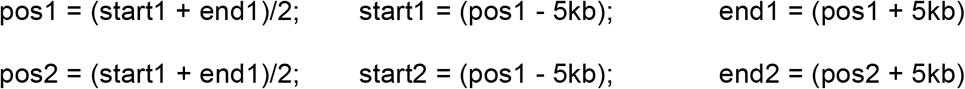

In order to overlap two lists, we performed 2 separate 1D overlaps with bedtools and then merged the results. To this end, every entry on each list is given a unique “loop ID.” Using *bedtools overlap* on each dimension of the loop list, we obtained a pair of loop IDs (one from each list) that were used to track which pairs of dots overlapped along both dimensions. Thus only pairs of dots with overlaps in both dimensions are merged and outputted.

#### Upset Plots

Upset plots were created for overlapping loops using the following R package: https://cran.r-project.org/web/packages/UpSetR/vignettes/basic.usage.html.

#### Quantification of chromatin loops

We created the loop pileups using notebooks from the *hic-data-analysis-bootcamp* notebook (https://github.com/hms-dbmi/hic-data-analysis-bootcamp/blob/master/notebooks/06_analysis_cooltools-snipping-pileups.ipynb). The pileups were done at 5kb resolution and with a 50kb extension on each side of the loop. To quantify the loop strength, first, we created an interaction matrix of 50×50 kb, centered around the loop. Then, we calculated the intensity of the loop by dividing the average of a 3×3 square in the middle of the interaction matrix by the average of its neighboring pixels; upper left, upper middle, upper right, lower left and right middle. See the image below:

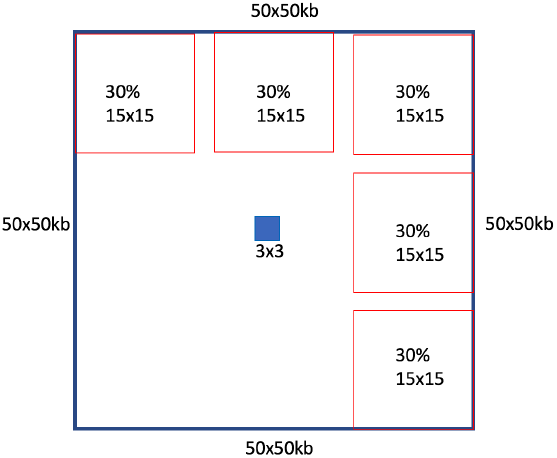

This quantified the loop enrichment using its local background, as was done to identify the loops. These quantifications are shown in figure 4b-4c, supplementary figure 4d-4e-4f.

#### Anchor Analysis

We concatenated the genomic positions of the left and the right anchors for each loop to create a 1D anchor list for each deep dataset (FA-DpnII, FA+DSG-DpnII, FA+DSG-MNase) derived from both H1-hESC and HFFc6 cell lines.

We used *BEDtools merge* (35) with “--c 1 -o count “ parameters to remove redundant anchors (based on their genomic position) and to find the number of merged anchors in each genomic location. The number of merged anchors in a given genomic locus reflected loop valency at this anchor. Using *BEDtools multiinter* (https://bedtools.readthedocs.io/en/latest/content/overview.html) we identified the anchors that were shared in 1,2 or 3 protocols (Figure 5a-5b-5c and Supplementary Figure 5a-5b-5c-5d-5e).

#### Cut&Run, Cut&Tag and ChIP Seq Analysis

Cut&Run data (HFFc6 H3K4me3, HFFc6 H3K27ac, H1-hESC CTCF, H1-hESC H3K4me3, H1-hESC H3K27ac) was generated in the lab of Steve Henikoff and can be found on the 4DN Data Portal (https://data.4dnucleome.org/). Cut&Tag (HFFc6 CTCF, HFFc6 SMC1) data was generated in the lab of René Maehr at UMass Medical school. Finally, ChIP Seq data was downloaded from ENCODE. We processed raw fastq files for Cut&Run and Cut&Tag data and downloaded already processed bigwig and peak lists for ChIP Seq data. We mapped and processed the fastq files using *nf-core ATAC Seq* (36) pipelines. BWA was used for mapping the fastq files to the hg38 reference genome; MACS2 (with default parameters) was used to find the enriched peaks and *BEDtools intersect* was subsequently used to identify the loop anchors from these enriched peaks.

We found the intersected anchors between the three protocols (FA-DpnII, FA+DSG-DpnII, FA+DSG-MNase) and the FA+DSG-MNase specific anchors using *bedtools intersect*. We extracted the open chromatin (ATAC Seq peaks) regions located at these anchors and then aggregated the average signal enrichments of CTCF, SMC1, H3K4me3, H3K27ac, YY1 and RNA PolII. *Deeptools* was used to create the enrichment profiles in Figure 5e and Supplementary Figure 5f (37). We downloaded the lists of candidate Cis Regulatory Elements (cCREs) for H1-hESC and HFFc6 from ENCODE (26) and overlapped these cCREs with the intersected anchor list and the FA+DSG-MNase anchor list, again using *BEDtools intersect*. Finally separated them based on the cCRE categories.

To compare the anchor specific enrichments shown in Fig 5g and Supplementary Figure 5h, we used the loop lists of FA-DpnII, FA+DSG-DpnII and FA+DSG-MNase. We identified enriched convergent CTCF sites located at these loop anchors and compared the enrichments of CTCF, SMC1, H3K4me3, H3K27ac, YY1 and RNA PolII per anchor. To obtain convergent CTCF sites, we selected Anchor 1 (left anchor) to overlap with CTCF sites that had a “+” orientation and a CTCF peak and Anchor 2 (right anchor) to overlap with CTCF sites that had a “-” orientation. We plotted convergent CTCF sites located at Anchor 1 and Anchor 2 for FA-DpnII, FA+DSG-DpnII and FA+DSG-MNase in both HFFc6 and H1-hESC (Figure 5f and Supplementary Fig. 5h).

For HFFc6, we used Cut&Tag data generated with an antibody against the N-terminus of CTCF. For H1-hESC cells, we used Cut&Run data generated with an antibody against the C-terminus of CTCF. Since CTCF motifs are known to locate at the N-terminus of the CTCF protein (27), the orientation of the CTCF enrichments differed between the data sets from Cut&Tag and Cut&Run.

#### Insulation Score

We calculated diamond insulation scores using cooltools (https://github.com/open2c/cooltools/blob/master/cooltools/cli/diamond_insulation.py) as implemented from Crane et al. (31) We defined the insulation and boundary strengths of each 10 kb bin by detecting the local minima of 10 kb binned data with a 200kb window size. We used cooltools’s *diamond-insulation* function with these parameters: “ --ignore-diags 2, --window-pixels 20”. We separated weak and strong boundaries using the mean insulation score of each protocol (i.e.: weak boundaries < mean < strong boundaries). Since diamond insulation pipelines cannot differentiate between compartment boundaries and insulation boundaries we manually removed the compartment boundaries before any further analysis. Therefore the depth in local minima here is a result of strong insulation strength not a compartment switch. Next, we aggregated the insulation strength of the deep datasets at loop anchors, strong boundaries and loop anchors located at the strong boundaries using scripts from the *hic-data-analysis-bootcamp* notebook (https://github.com/hms-dbmi/hic-data-analysis-bootcamp/blob/master/notebooks/06_analysis_cooltools-snipping-pileups.ipynb). For both deep and matrix data we used only strong boundaries for further analysis since they reflected the true boundaries across protocols. Since the position of insulation boundaries was often offset by one or two bins between protocols, we extended the boundary bin by 10 kb on each side (30 kb total) in each protocol. We then used *bedtools multiinter* (https://bedtools.readthedocs.io/en/latest/content/overview.html) to count the boundaries that were found in one or more protocols within the cell type. We defined our stringent boundary list as the boundaries that were shared in at least 50% of the matrix protocols within each cell type and used these boundary lists for further comparisons. In heatmaps, we used the average insulation strength of these boundaries per protocol (Supplementary Fig 6k). To create the heatmaps in Supplementary Fig. 6l, we used the loop anchors that were shared between the 3 protocols that were deeply sequenced: FA+DpnII, FA+DSG-DpnII and FA+DSG-MNase in both H1-hESC and HFFc6.

#### Loop quantification for specific genomic separations

To quantify the loop strengths for HFFc6 deep datasets described in Fig 6d (FA-DpnII, FA+DSG-DpnII, FA+DSG-DdeI, FA+DSG-DdeI-DpnII, FA+DSG-MNase), first, we seperated the loops based on their genomic separations into 100 kb bins, starting from 70kb (i.e. 70-170-kb, 170-270kb,…970-1070 kb), because 70kb was the smallest detectable loop size and then plotted the number of loops detected in each distance interval (Fig. 6d). Since the number of detected loops in these genomic separations was different for each library, we sampled 1,000 loops for each distance from FA+DSG-DdeI-DpnII to quantify loop enrichments of the 5 libraries (Fig. 6e). If the number of loops at a specified distance was smaller than 1000 we use the entire loop set at this distance.

Finally, to create Fig. 6h and Fig. 6i we sampled 2,000 loops from each HFFc6 deep dataset, (FA-DpnII, FA+DSG-DpnII, FA+DSG-DdeI, FA+DSG-DdeI-DpnII, FA+DSG-MNase), combined them and then quantified the loop strength of the total 10,000 loops in these deep datasets (Fig. 6h) and in matrix datasets described in Fig. 1a (Fig. 6i). Loop enrichments were quantified as described in the “Quantification of chromatin loops” section.

