## Supplemental Figures for "Systematic evaluation of chromosome conformation capture assays"

### Supplemental Material – Table of contents

#### Supplementary Figures

Supplemental figure 1 - DNA fragmentation and hierarchical clustering of distance corrected correlation (HiCRep)

Supplemental figure 2 - Distance dependent interaction frequency and the number of inter-chromosomal interactions change for data generated with protocols that use various enzyme and cross-linker combinations

Supplemental figure 3 - Compartment identity is robust for protocol variation but the strength of the compartments differs between protocols

Supplemental figure 4 - Chromatin loops are better detected in experiments with fine fragmentation and DSG cross-linking

Supplemental figure 5 - The number of detectable loops associated within a given anchor increases with extra cross-linking and smaller fragmentation. This also enhances the detectability of CTCF depleted promoter enhancer loops

Supplemental figure 6 - Insulation boundaries show modest differences across experimental variations

Supplemental figure 7 - Experiments performed using two enzymes have smaller fragments

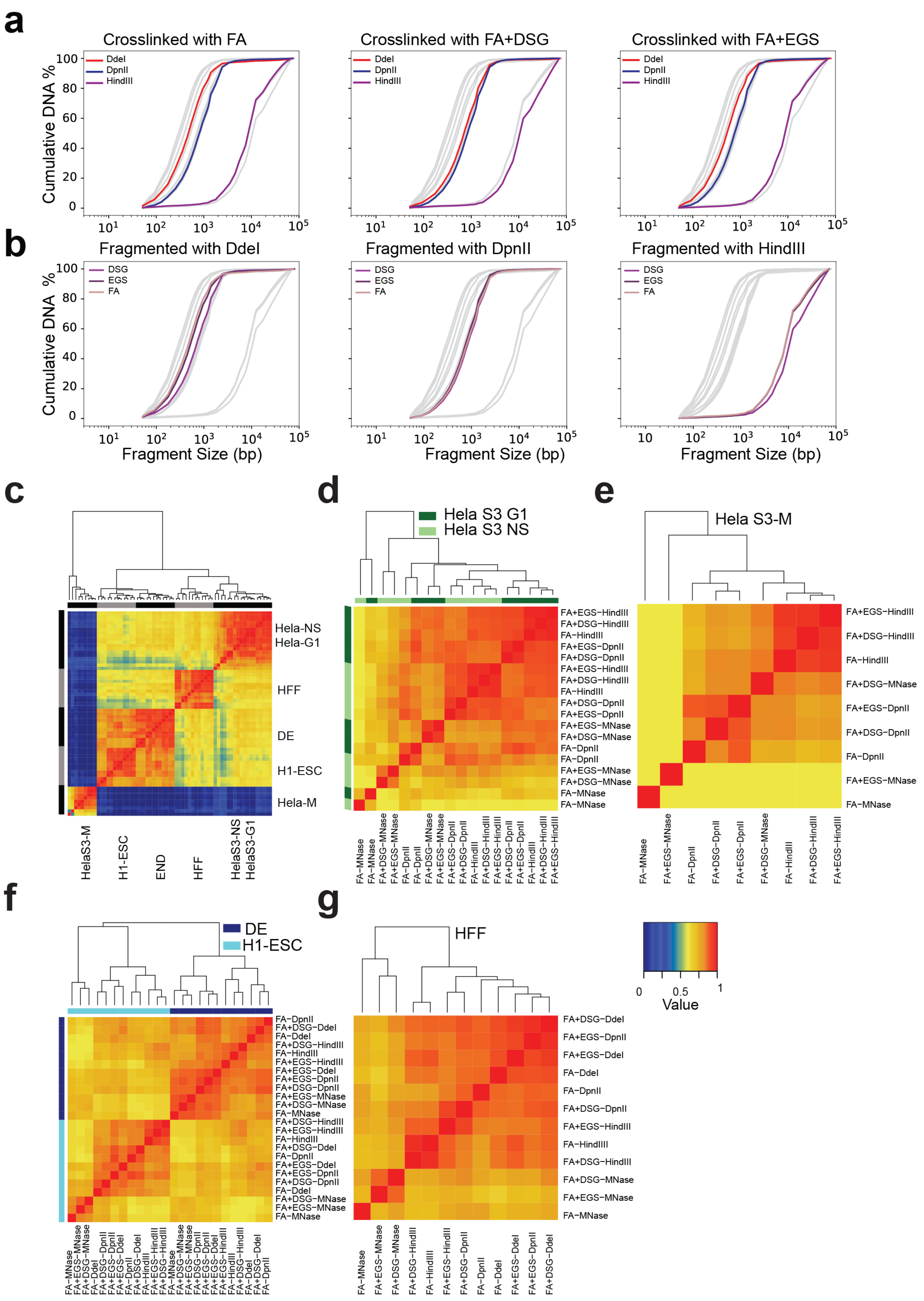
a
Crosslinked with FA
Crosslinked with FA+DSG
Crosslinked with FA+EGS
DdeI
DpnII
HindIII
Cumulative DNA %
b
Fragmented with DdeI
Fragmented with DpnII
Fragmented with HindIII
DSG
EGS
FA
Fragment Size (bp)
c
Hela-NS
Hela-G1
HFF
DE
H1-ESC
Hela-M
HelaS3-M
H1-ESC
END
HFF
HelaS3-NS
HelaS3-G1
d
Hela S3 G1
Hela S3 NS
e
Hela S3-M
f
DE
H1-ESC
g
HFF
Value

**Supplementary Fig. 1: DNA fragmentation and hierarchical clustering of distance corrected correlation (HiCRep)**

a,b Cumulative distribution of the lengths of fragmented DNA obtained from fragment analyzer data in HFF cells stratified for different cross-linkers (a) and restriction enzymes (b). Gray lines indicate all datasets, colored lines indicate data obtained with the indicated nuclease/crosslinkers.

c-g Hierarchical clustering of HiCRep correlations for: all protocols comparing cell states (c), synchronized Hela S3 G1 cells (dark green) and non-synchronized Hela S3 cells (light green) (d), synchronized Hela S3 mitotic cells (e), H1-hESC and H1-hESC derived DE cells (f), 12 protocols applied to HFF cells (g). One color key is indicated for all of the heatmaps.

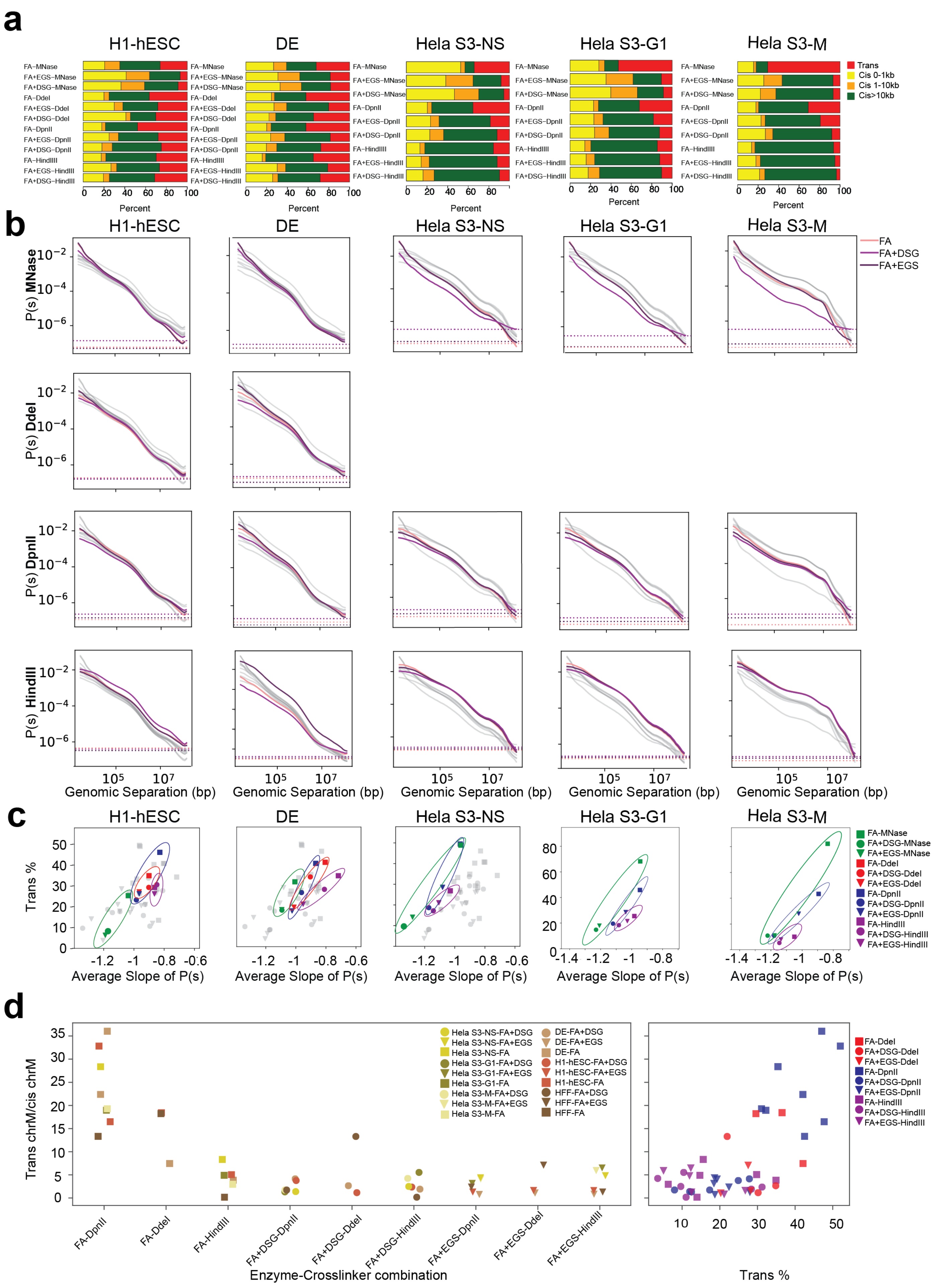

a
H1-hESC
DE
Hela S3-NS
Hela S3-G1
Hela S3-M
FA−MNase
FA+EGS−MNase
FA+DSG−MNase
FA−DdeI
FA+EGS−DdeI
FA+DSG−DdeI
FA−DpnII
FA+EGS−DpnII
FA+DSG−DpnII
FA−HindIIII
FA+EGS−HindIII
FA+DSG−HindIII
Trans
Cis 0-1kb
Cis 1-10kb
Cis>10kb
Percent
b
P(s) MNase
P(s) DdeI
P(s) DpnII
P(s) HindIII
FA
FA+DSG
FA+EGS
Genomic Separation (bp)
c
Trans %
Average Slope of P(s)
FA-MNase
FA+DSG-MNase
FA+EGS-MNase
FA-DdeI
FA+DSG-DdeI
FA+EGS-DdeI
FA-DpnII
FA+DSG-DpnII
FA+EGS-DpnII
FA-HindIII
FA+DSG-HindIII
FA+EGS-HindIII
d
Trans chrM/cis chrM
Hela S3-NS-FA+DSG
Hela S3-NS-FA+EGS
Hela S3-NS-FA
Hela S3-G1-FA+DSG
Hela S3-G1-FA+EGS
Hela S3-G1-FA
Hela S3-M-FA+DSG
Hela S3-M-FA+EGS
Hela S3-M-FA
DE-FA+DSG
DE-FA+EGS
DE-FA
H1-hESC-FA+DSG
H1-hESC-FA+EGS
H1-hESC-FA
HFF-FA+DSG
HFF-FA+EGS
HFF-FA
FA-DpnII
FA-DdeI
FA-HindIII
FA+DSG-DpnII
FA+DSG-DdeI
FA+DSG-HindIII
FA+EGS-DpnII
FA+EGS-DdeI
FA+EGS-HindIII
Enzyme-Crosslinker combination
FA-DdeI
FA+DSG-DdeI
FA+EGS-DdeI
FA-DpnII
FA+DSG-DpnII
FA+EGS-DpnII
FA-HindIII
FA+DSG-HindIII
FA+EGS-HindIII
Trans %

**Supplementary Fig. 2: Distance dependent interaction frequency and the number of inter-chromosomal interactions change for data generated with protocols that use various enzyme and cross-linker combinations**.

a The number of valid pairs in each of the 12 protocols applied to H1-hESC, DE, Hela S3-NS, Hela S3-G1 and Hela S3-M cells partitioned by genomic distances

b Distance dependent contact probability of 12 protocols ordered as in (a), partitioned by fragmenting nuclease used (gray lines indicate all datasets, colored lines indicate datasets generated with the nucleases indicated for each plot).

c The relationship between the trans percent and the average slope of the distance dependent contact probability for the 12 protocols ordered as in (a).

d Quantification of protocol introduced noise as defined by inter-mitochondrial interactions (chrM with chr1-22), normalized by intra-mitochondrial (chrM with chrM) interactions.

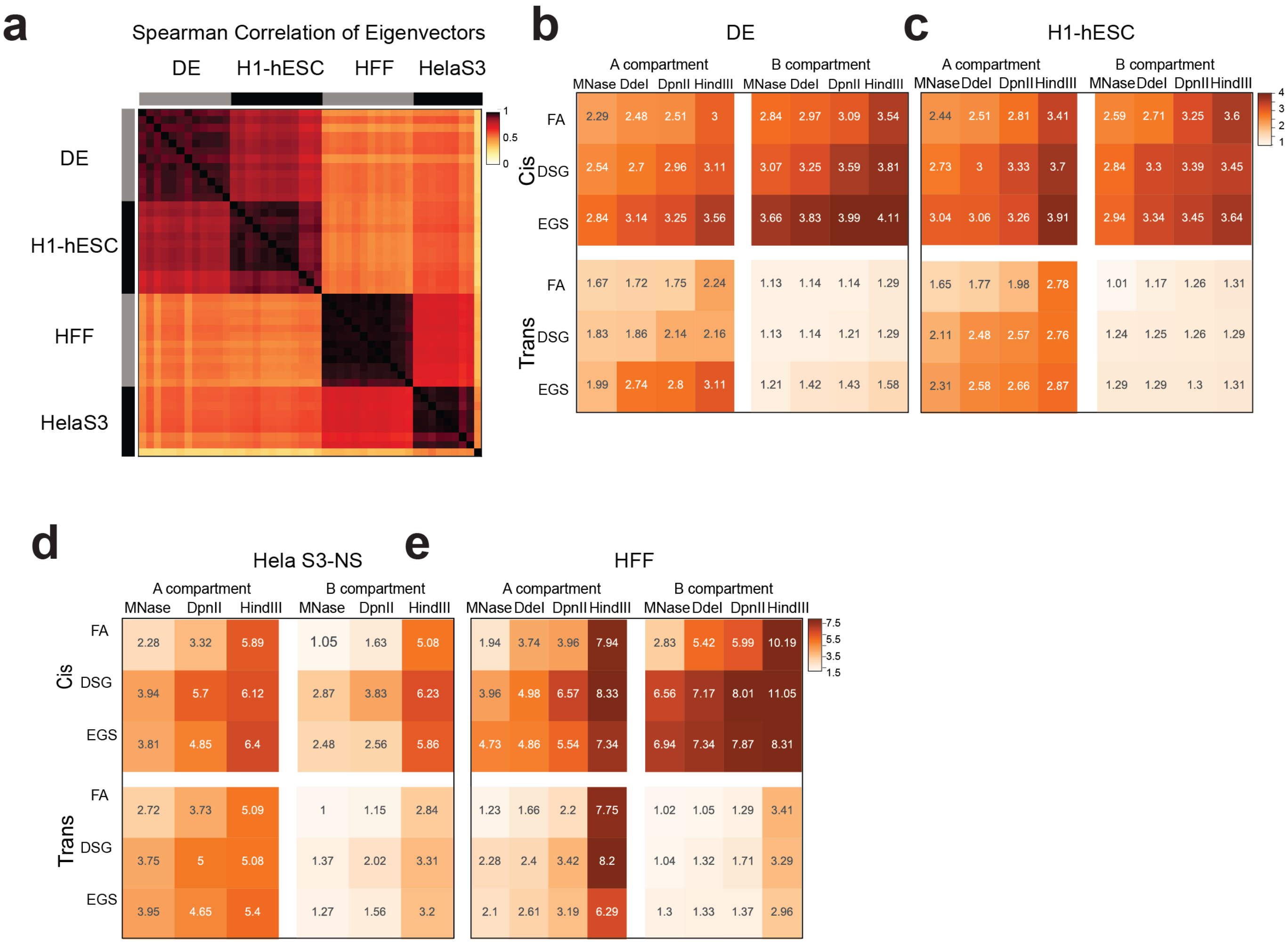


**Supplemental Fig. 3: Compartment identity is robust for protocol variation but the strength of the compartments differs between protocols**

a Hierarchical clustering of Spearman correlations of Eigenvectors (PC1) for 63 protocols. Clustering shows strong correlations between compartments (PC1 values) from data obtained with varying protocols applied to the same cell types and weaker correlations for data obtained with the same protocols applied to different cell types.

b-e A-A and B-B compartment strength of saddle plots for fixation versus enzyme stratified by cell state: DE (b), H1-hESC (c), Hela S3-NS (d), HFF (e). For each cell type, saddle plot quantification was done for cis and trans reads separately.

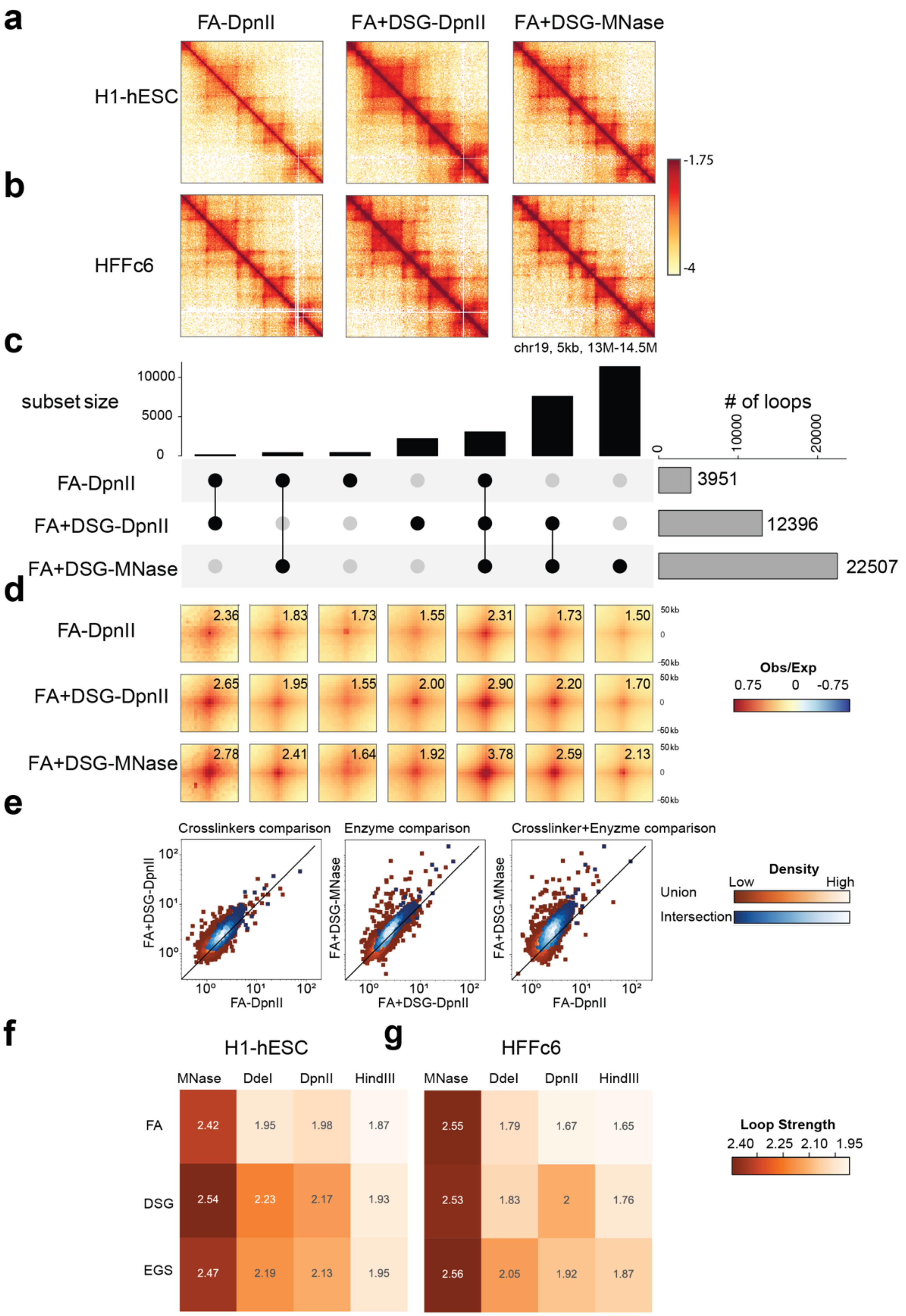

a
b
c
d
e
f
g
FA-DpnII
FA+DSG-DpnII
FA+DSG-MNase
H1-hESC
HFFc6
-1.75
-4
chr19, 5kb, 13M-14.5M
subset size
10000
5000
0
### of loops
0
10000
20000
3951
12396
22507
2.36 1.83 1.73 1.55 2.31 1.73 1.50
2.65 1.95 1.55 2.00 2.90 2.20 1.70
2.78 2.41 1.64 1.92 3.78 2.59 2.13
50kb
0
-50kb
Obs/Exp
0.75 0 -0.75
Crosslinkers comparison
Enzyme comparison
Crosslinker+Enyzme comparison
FA+DSG-DpnII
FA+DSG-MNase
FA-DpnII
FA+DSG-DpnII
10⁰ 10¹ 10²
Density
Low High
Union
Intersection
H1-hESC
HFFc6
MNase DdeI DpnII HindIII
FA 2.42 1.95 1.98 1.87
DSG 2.54 2.23 2.17 1.93
EGS 2.47 2.19 2.13 1.95
FA 2.55 1.79 1.67 1.65
DSG 2.53 1.83 2 1.76
EGS 2.56 2.05 1.92 1.87
Loop Strength
2.40 2.25 2.10 1.95

**Supplementary Fig. 4: Chromatin loops are better detected in experiments with fine fragmentation and DSG cross-linking**

a Interaction heatmaps of experiments for H1-ESC cells obtained from the following crosslinker-enzyme combinations (from left to right): FA-DpnII, FA+DSG-DpnII and FA+DSG-MNase.

b Interaction heatmaps of protocols specified in Supplementary Fig. 4a for HFFc6 cells.

c Upset plot of loops detected in protocols performed in H1-hESC showing 1) total number of loops detected in FA-DpnII, FA+DSG-DpnII and FA+DSG-MNase (gray bars on the right), 2) number of loops detected in one, two or three of these protocols shown as black bars. The combinations highlighted with connected black dots.

d Pileups of the loops detected in all combinations for H1-hESC protocols shown in Fig 4a. Quantification of the loop strength was done taking an observed/expected interaction matrix (50x50 kb) around the loop and then normalizing the loop intensity to its local background (see methods).

e Scatter plots with relative strengths of individual loops between pairs of protocols for H1-hESC; from left to right: FA-DpnII v/s FA+DSG-DpnII (different crosslinking with the same enzyme), FA+DSG-DpnII v/s FA+DSG-MNase (different enzymes with same crosslinking) and FA-DpnII v/s FA+DSG-MNase (different enzymes and different crosslinking). Loop strengths were calculated as in panel d but for individual looping interactions. Scatter plots were drawn for two sets of looping interactions (1) the union of all locations from the three protocols (red squares) and (2) interaction of all locations from the three protocols (blue circles). Color scale represents the density of looping interactions.

f,g Quantification of aggregated loop strengths from the matrix of 12 protocols described in Fig 1a for H1-hESC cells (f), and HFFc6 cell (g). Pileups represent looping interactions detected across all three deep protocols (FA-DpnII, FA+DSG-DpnII and FA+DSG-MNase) in each cell type

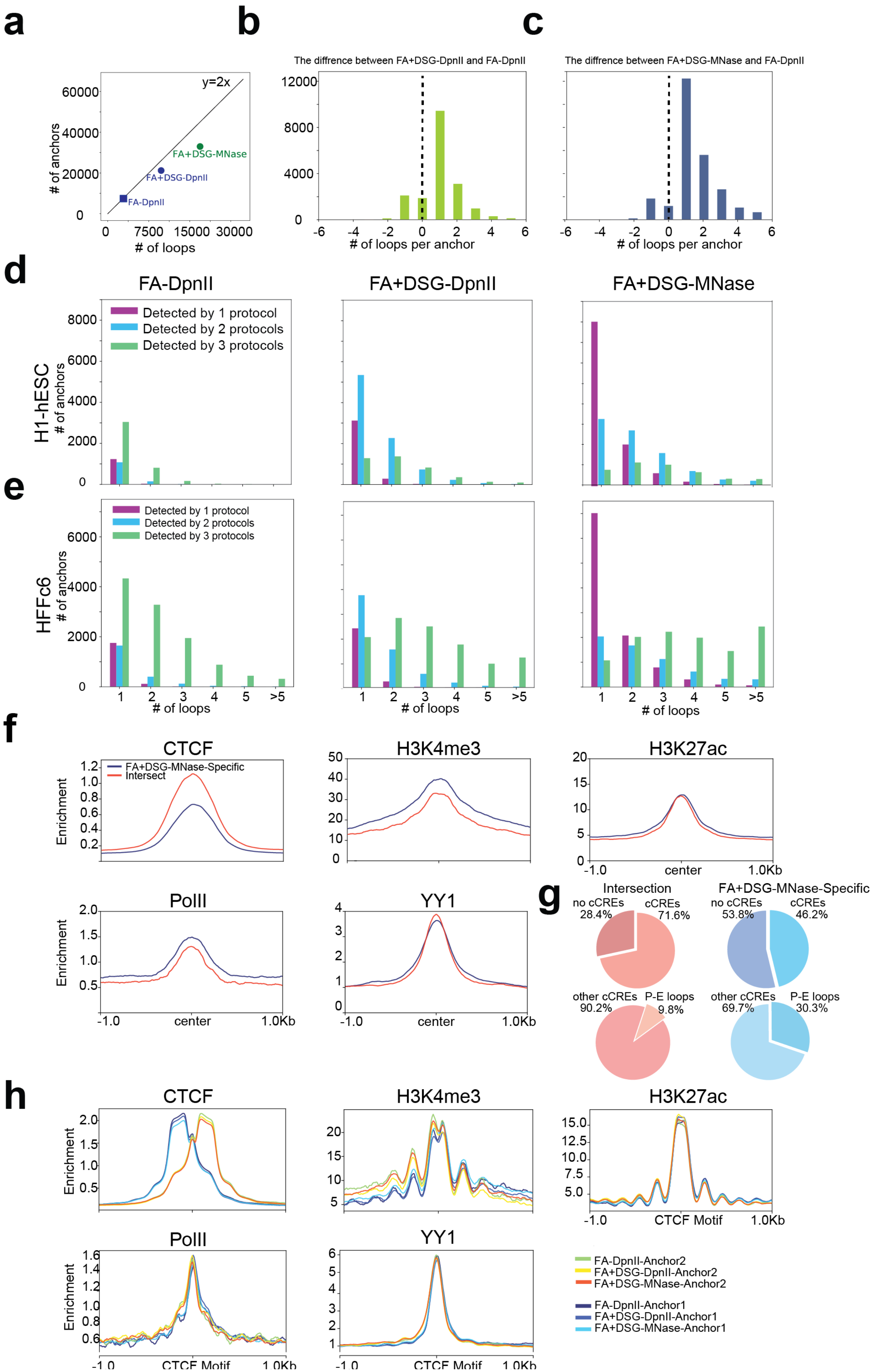

a
b
c
y=2x
FA+DSG-MNase
FA+DSG-DpnII
FA-DpnII
### of anchors
### of loops
The diffrence between FA+DSG-DpnII and FA-DpnII
The diffrence between FA+DSG-MNase and FA-DpnII
### of loops per anchor
d
FA-DpnII
FA+DSG-DpnII
FA+DSG-MNase
Detected by 1 protocol
Detected by 2 protocols
Detected by 3 protocols
H1-hESC
### of anchors
e
HFFc6
### of anchors
### of loops
f
CTCF
H3K4me3
H3K27ac
FA+DSG-MNase-Specific
Intersect
Enrichment
PolII
YY1
-1.0
center
1.0Kb
g
Intersection
FA+DSG-MNase-Specific
no cCREs 28.4%
cCREs 71.6%
no cCREs 53.8%
cCREs 46.2%
other cCREs 90.2%
P-E loops 9.8%
other cCREs 69.7%
P-E loops 30.3%
h
CTCF
H3K4me3
H3K27ac
PolII
YY1
Enrichment
CTCF Motif
FA-DpnII-Anchor2
FA+DSG-DpnII-Anchor2
FA+DSG-MNase-Anchor2
FA-DpnII-Anchor1
FA+DSG-DpnII-Anchor1
FA+DSG-MNase-Anchor1

**Supplementary Fig. 5: The number of detectable loops associated within a given anchor increases with extra cross-linking and smaller fragmentation. This also enhances the detectability of CTCF depleted promoter enhancer loops.**

a The number of loops detected in H1-hESC (x-axis) plotted against the number of loop anchors (y-axis). y=2x line shows the expected relationship between loops and loop anchors when each anchor is engaged in only 1 loop.

b,c The number of FA-DpnII loops subtracted from the number of FA+DSG-DpnII loops (b) or the number of FA+DSG-MNase loops (c) detected at the same anchors. The union of loop calls from both of the plotted protocols was used here.

d Histograms of valencies of loop anchors detected in H1-hESC. Each panel represents loop anchors found in the given protocol (from left to right, FA-DpnII, FA+DSG-DpnII, FA+DSG-MNase). For each protocol, the anchors were further stratified into three categories. The leftmost panel (FA-DpnII) is used as a guiding example. The categories are: anchors detected by 1 protocol (FA-DpnII), anchors detected in 2 protocols (FA-DpnII and either FA+DSG-DpnII or FA+DSG-MNase) and anchors detected in all 3 protocols (FA-DpnII and FA+DSG-DpnII and FA+DSG-MNase ).

e The same plots as shown in Supplementary Fig. 5d but generated using HFFc6 cells.

f The comparison of CTCF, H3K4me3, H3K27ac, PolII, and YY1 enrichments at loop anchors centered at open chromatin regions. Open chromatin regions (as quantified by ATAC Seq) that are located within the anchor coordinates are used to center the average enrichments. Anchors detected by all protocols and FA+DSG-MNase-specific anchors in H1-hESC.

g Top row: candidate Cis Regulatory elements (cCREs) in common (left) and FA+DSG-MNase specific loop anchors (right), as specified in Supplemantary Figure 5f. Bottom row: the percentage cCREs for Promoter-Enhancer elements without CTCF enrichment.

h The enrichment of CTCF, SMC1, H3K4me3 and H3K27ac in left (Anchor1) and right (Anchor 2) anchors centered by CTCF sites. Loop anchors present in FA-DpnII, FA+DSG-DpnII or FA+DSG-MNase applied to HFFc6.

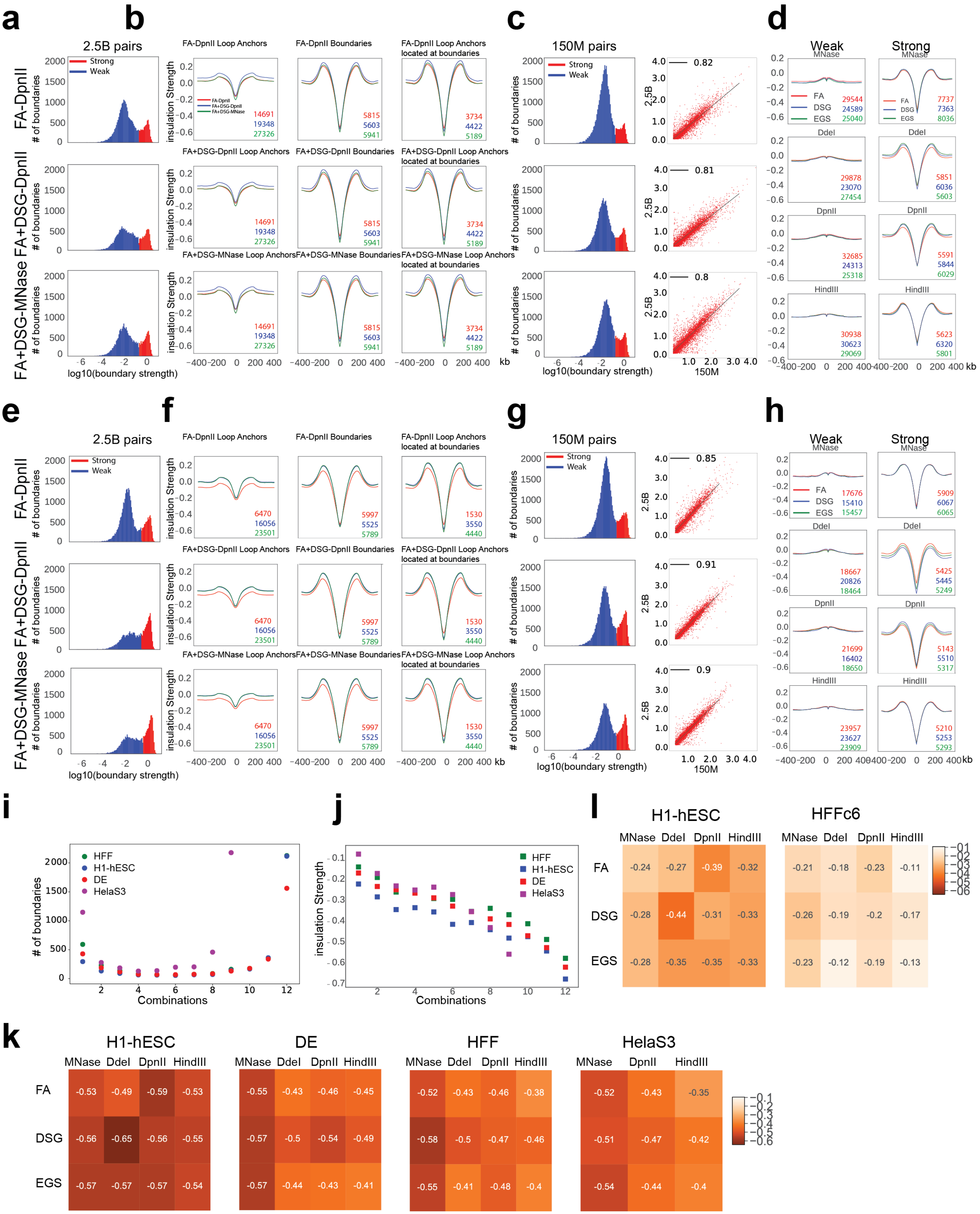

a
b
c
d
2.5B pairs
Strong
Weak
FA-DpnII
FA+DSG-DpnII
FA+DSG-MNase
### of boundaries
log10(boundary strength)
FA-DpnII Loop Anchors
FA-DpnII Boundaries
FA-DpnII Loop Anchors located at boundaries
FA+DSG-DpnII Loop Anchors
FA+DSG-DpnII Boundaries
FA+DSG-DpnII Loop Anchors located at boundaries
FA+DSG-MNase Loop Anchors
FA+DSG-MNase Boundaries
FA+DSG-MNase Loop Anchors located at boundaries
insulation Strength
14691
19348
27326
5815
5603
5941
3734
4422
5189
kb
150M pairs
2.5B
150M
0.82
0.81
0.8
Weak
Strong
MNase
DdeI
DpnII
HindIII
FA
DSG
EGS
29544
24589
25040
7737
7363
8036
29878
23070
27454
5851
6036
5603
32685
24313
25318
5591
5844
6029
30938
30623
29069
5623
6320
5801
e
f
g
h
6470
16056
23501
5997
5525
5789
1530
3550
4440
0.85
0.91
0.9
17676
15410
15457
5909
6067
6065
18667
20826
18464
5425
5445
5249
21699
16402
18650
5143
5510
5317
23957
23627
23909
5210
5253
5293
i
HFF
H1-hESC
DE
HelaS3
### of boundaries
Combinations
j
insulation Strength
Combinations
l
H1-hESC
HFFc6
MNase DdeI DpnII HindIII
FA -0.24 -0.27 -0.39 -0.32
DSG -0.28 -0.44 -0.31 -0.33
EGS -0.28 -0.35 -0.35 -0.33
FA -0.21 -0.18 -0.23 -0.11
DSG -0.26 -0.19 -0.2 -0.17
EGS -0.23 -0.12 -0.19 -0.13
k
H1-hESC
MNase DdeI DpnII HindIII
FA -0.53 -0.49 -0.59 -0.53
DSG -0.56 -0.65 -0.56 -0.55
EGS -0.57 -0.57 -0.57 -0.54
DE
MNase DdeI DpnII HindIII
FA -0.55 -0.43 -0.46 -0.45
DSG -0.57 -0.5 -0.54 -0.49
EGS -0.57 -0.44 -0.43 -0.41
HFF
MNase DdeI DpnII HindIII
FA -0.52 -0.43 -0.46 -0.38
DSG -0.58 -0.5 -0.47 -0.46
EGS -0.55 -0.41 -0.48 -0.4
HelaS3
MNase DpnII HindIII
FA -0.52 -0.43 -0.35
DSG -0.51 -0.47 -0.42
EGS -0.54 -0.44 -0.4

**Supplementary Fig. 6: Insulation boundaries show modest differences across experimental variations**

a Boundary strength distribution (log scale) for HFFc6 deep data (~2.5 B valid pairs) from FA-DpnII (top) , FA+DSG-DpnII (middle) and FA+DSG-MNase (bottom). Boundaries are classified as weak (blue) or strong (red) based on boundary strength (see methods)

b Pileups in FA-DpnII (top row) , FA+DSG-DpnII (middle row) and FA+DSG-MNase (bottom row) for aggregate insulation scores at loop anchors (left), strong insulation boundaries (middle) and loop anchors colocalizing with strong insulation boundaries (right) as detected in deeply sequenced libraries: FA-DpnII (red), FA+DSG-DpnII (blue) and FA+DSG-MNase (green).

c The effect of sequencing depth on boundary strength. Left panel shows the boundary strength distribution of matrix data (Fig. 1a, ~150 M valid pairs) for FA-DpnII (top) , FA+DSG-DpnII (middle) and FA+DSG-MNase (bottom) applied to HFFc6 cells. An excess of weak boundaries is observed when comparing to the equivalent deeply sequenced library as shown in Supplementary Fig. 6a. Focusing on the strong boundaries, the right panel shows a strong correlation between deep and matrix data for FA-DpnII, FA+DSG-DpnII and FA+DSG-MNase.

d Aggregate insulation profile of the boundaries obtained from matrix data for HFFc6 stratified by crosslinkers and nucleases. Boundaries further separated based on their insulation strength as weak and strong insulation.

e-h H1-hESC data displayed like Supplementary Fig. 6a-d.

i Distributions of the number of boundaries (y-axis) stratified by the number of protocols in which a given boundary was detected (x-axis). The number of protocols varies between 1 to 12 (Fig 1a).

j Insulation strength of the boundaries stratified in the same manner as Supplementary Fig 6i.

k Mean insulation strength from boundaries detected in at least half of the protocols for various cross-linkers and enzyme combinations of H1-hESC, DE, HFF and Hela S3-NS (see methods).

l Mean insulation strength of loop anchors that are detected in all three deep protocols (FA-DpnII, FA+DSG-DpnII and FA+DSG-MNase) for both HFFc6 and H1-hESC, averaged for 12 protocols of H1-hESC and HFF.

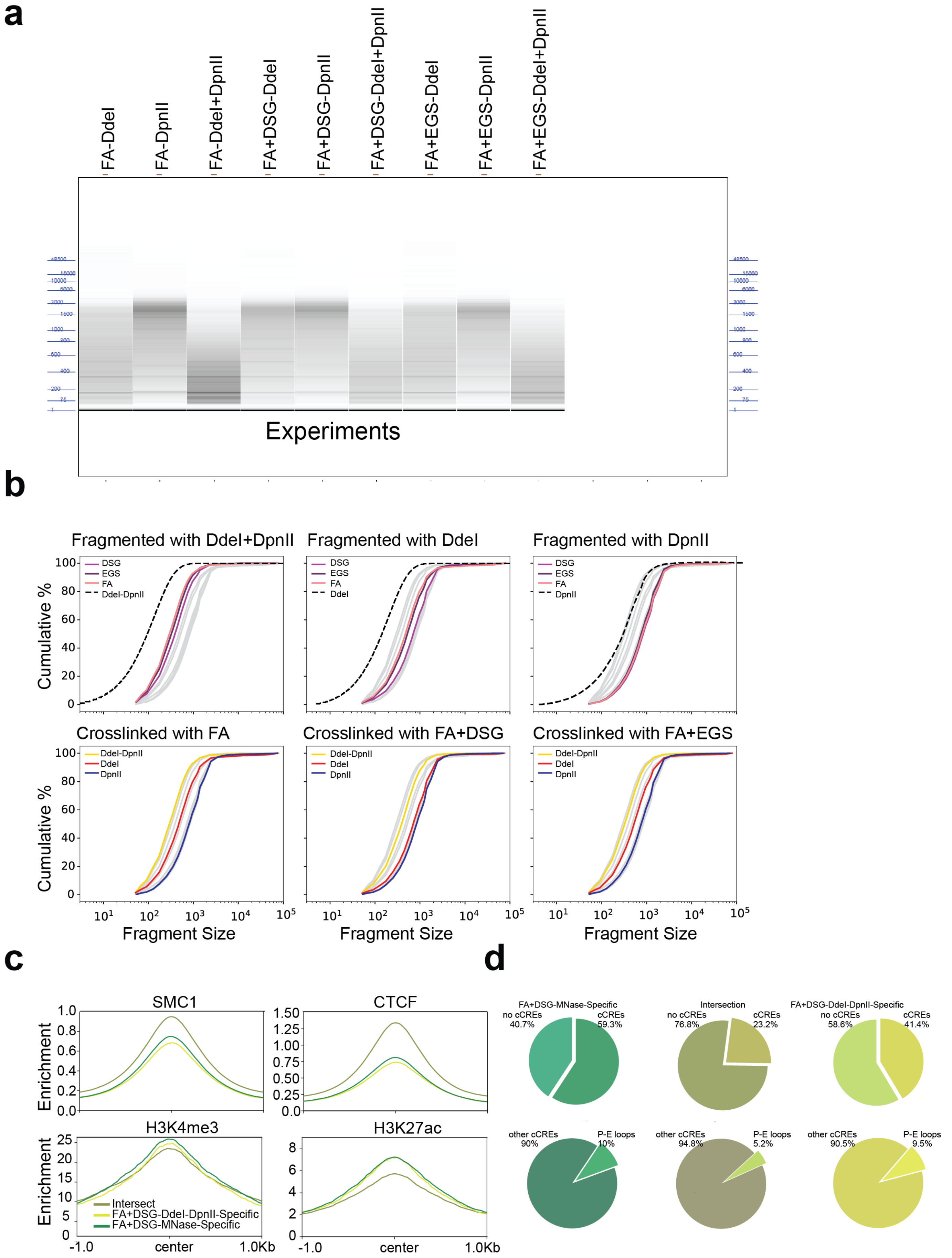

a
FA-DdeI
FA-DpnII
FA-DdeI+DpnII
FA+DSG-DdeI
FA+DSG-DpnII
FA+DSG-DdeI+DpnII
FA+EGS-DdeI
FA+EGS-DpnII
FA+EGS-DdeI+DpnII
48500
15000
10000
6000
3000
1500
1000
800
600
400
200
75
1
Experiments
b
Fragmented with DdeI+DpnII
Fragmented with DdeI
Fragmented with DpnII
DSG
EGS
FA
DdeI-DpnII
DdeI
DpnII
Cumulative %
Crosslinked with FA
Crosslinked with FA+DSG
Crosslinked with FA+EGS
Fragment Size
c
SMC1
CTCF
H3K4me3
H3K27ac
Enrichment
Intersect
FA+DSG-DdeI-DpnII-Specific
FA+DSG-MNase-Specific
-1.0
center
1.0Kb
d
FA+DSG-MNase-Specific
no cCREs 40.7%
cCREs 59.3%
Intersection
no cCREs 76.8%
cCREs 23.2%
FA+DSG-DdeI-DpnII-Specific
no cCREs 58.6%
cCREs 41.4%
other cCREs 90%
P-E loops 10%
other cCREs 94.8%
P-E loops 5.2%
other cCREs 90.5%
P-E loops 9.5%

**Supplementary Fig. 7: Experiments performed using two enzymes have smaller fragments.**

a Fragment size distributions from Fragment Analyzer for specified protocols.

b Cumulative distributions of fragmented DNA in HFF cells stratified for cross-linking agents (top row) or restriction enzymes (bottom row). Dashed lines in each of the panels represent expected fragment size distribution from *in silico* digestion of hg38 for enzymes indicated. Gray lines represent all data from all other enzymes (columns).

c Comparison of CTCF, SMC1, H3K4me3 and H3K27ac enrichments at loop anchors centered at open chromatin regions. Open chromatin regions (ATAC Seq) located within the anchor coordinates were used to center the average enrichments. Anchors were separated into sets detected by FA+DSG-DdeI+DpnII, FA+DSG-MNase or both.

d Percentage of cCREs and promoter-enhancer elements located at loop anchors specific to FA+DSG-DdeI+DpnII, FA+DSG-MNase or shared between them.
